# A cysteine protease inhibitor blocks SARS-CoV-2 infection of human and monkey cells

**DOI:** 10.1101/2020.10.23.347534

**Authors:** Drake M. Mellott, Chien-Te Tseng, Aleksandra Drelich, Pavla Fajtová, Bala C. Chenna, Demetrios H. Kostomiris, Jason Hsu, Jiyun Zhu, Zane W. Taylor, Vivian Tat, Ardala Katzfuss, Linfeng Li, Miriam A. Giardini, Danielle Skinner, Ken Hirata, Sungjun Beck, Aaron F. Carlin, Alex E. Clark, Laura Beretta, Daniel Maneval, Felix Frueh, Brett L. Hurst, Hong Wang, Klaudia I. Kocurek, Frank M. Raushel, Anthony J. O’Donoghue, Jair Lage de Siqueira-Neto, Thomas D. Meek, James H. McKerrow

**Affiliations:** Departments of Biochemistry and Biophysics, Texas A&M University, 301 Old Main Drive, College Station, Texas 77843; Department of Chemistry, Texas A&M University, 301 Old Main Drive, College Station, Texas 77843; Department of Microbiology and Immunology, University of Texas, Medical Branch, 3000 University Boulevard, Galveston, Texas, 77755-1001; Skaggs School of Pharmacy and Pharmaceutical Sciences, University of California San Diego, La Jolla, CA; Institute of Organic Chemistry and Biochemistry, Academy of Sciences of the Czech Republic, 16610 Prague, Czech Republic; Selva Therapeutics, 5600 Old Main Hill, Utah State University, Logan, Utah, 84322; Institute for Antiviral Research, Department of Animal, Dairy, and Veterinary Sciences, 5600 Old Main Hill, Utah State University, Logan, Utah, 84322; Department of Medicine, Division of Infectious Diseases and Global Public Health, University of California, San Diego, La Jolla, CA 92037, USA; Biological Sciences Division, Pacific Northwest National Laboratory, 902 Battelle Blvd, Richland, WA 99353

**Author notes:** Thomas D. Meek and James H. McKerrow are co-Senior Authors. **CONTRIBUTIONS** Experimental design was provided by J.H.M., D.M.M., C.H.T., J.L.S-N., A.D, T.D.M., F.F., A.J.O., Z.W.T. and F.M.R. These and other authors provided experimental data. The manuscript was written by J.H.M., D.M.M., C.H.T., A.J.O., J.L.N-S., F.F, and T.D.M.

**Keywords:** SARS-CoV-2, protease inhibitor, K777, cathepsin L, spike protein

## Abstract

K777 is a di-peptide analog that contains an electrophilic vinyl-sulfone moiety and is a potent, covalent inactivator of cathepsins. Vero E6, HeLa/ACE2, Caco-2, A549/ACE2, and Calu-3, cells were exposed to SARS-CoV-2, and then treated with K777. K777 reduced viral infectivity with EC_50_ values of inhibition of viral infection of: 74 nM for Vero E6, <80 nM for A549/ACE2, and 4 nM for HeLa/ACE2 cells. In contrast, Calu-3 and Caco-2 cells had EC_50_ values in the low micromolar range. No toxicity of K777 was observed for any of the host cells at 10-100 μM inhibitor. K777 did not inhibit activity of the papain-like cysteine protease and 3CL cysteine protease, encoded by SARS-CoV-2 at concentrations of ≤ 100 μM. These results suggested that K777 exerts its potent anti-viral activity by inactivation of mammalian cysteine proteases which are essential to viral infectivity. Using a propargyl derivative of K777 as an activity-based probe, K777 selectively targeted cathepsin B and cathepsin L in Vero E6 cells. However only cathepsin L cleaved the SARS-CoV-2 spike protein and K777 blocked this proteolysis. The site of spike protein cleavage by cathepsin L was in the S1 domain of SARS-CoV-2, differing from the cleavage site observed in the SARS CoV-1 spike protein. These data support the hypothesis that the antiviral activity of K777 is mediated through inhibition of the activity of host cathepsin L and subsequent loss of viral spike protein processing.

**SIGNIFICANCE:** The virus causing COVID-19 is highly infectious and has resulted in a global pandemic. We confirm that a cysteine protease inhibitor, approved by the FDA as a clinical-stage compound, inhibits SARS-CoV-2 infection of several human and monkey cell lines with notable(nanomolar) efficacy. The mechanism of action of this inhibitor is identified as a specific inhibition of host cell cathepsin L. This in turn inhibits host cell processing of the coronaviral spike protein, a step required for cell entry. Neither of the coronaviral proteases are inhibited, and the cleavage site of spike protein processing is different from that reported in other coronaviruses. Hypotheses to explain the differential activity of the inhibitor with different cell types are discussed.

## INTRODUCTION

The COVID-19 outbreak of 2019(1,2) has led to a global health crisis of a magnitude not seen since the influenza pandemic of 1918. By October 2020, more than 40 million people have been infected worldwide, including more than 8 million in the US(3, 4). The scope of COVID-19 pandemic is such that as-yet unavailable vaccines may be too late to circumvent a prolonged health and economic crisis. Accordingly, there is an urgent need for the rapid identification of therapies that limit the pathology caused by SARS-CoV-2 for the management of COVID-19(5,6).

The severe acute respiratory syndrome (SARS) and Middle East respiratory syndrome (MERS) outbreaks of 2003 and 2012(7,8), caused by betacoronaviruses highly related to SARS CoV-2 provided important clues to the mechanism of viral infection of host cells. These include two virally-encoded cysteine proteases, 3CL protease (Mpro)(10) and a papain-like protease (PLpro)(11) that are essential to coronaviral maturation. In addition, the trimeric coronaviral spike glycoprotein of SARS-CoV-2 (76% sequence identity to SARS-CoV-1 spike protein (12)), is involved in the fusion of the viral envelope with mammalian cell membranes, followed by the release of the genomic viral RNA and the nucleocapsid complex into the host cell (cellular entry)(13, 14). Studies of the SARS-CoV-1 spike protein have shown that it adheres to the extracellular domain of the angiotensin converting enzyme-2 (ACE2) receptor in susceptible human cells (12, 15–16). In SARS-CoV-2, the receptor binding domain of the spike protein subunit S1 binds to the ACE2 receptor, and the subunit S2 is involved in the viral cell membrane fusion to facilitate viral entry (18–19). Prior to cellular entry, the spike protein requires processing by host cell proteases for activation (16–18, 21). In particular, the transmembrane protease serine-2 (TMPRSS-2), the cysteine protease cathepsin L (CTSL), and furin-like proteases, have been proposed to play roles in preparing SARS-CoV-2 to enter cells by either endocytosis or traversing the cellular membrane (18–22).

These proposed mechanisms have stimulated studies to explore the role of both serine and cysteine protease inhibitors in reducing coronaviral entry. Inhibitors of TMPRSS-2, such as camostat, reduced infectivity in selected human cell lines (18). Broad spectrum cysteine protease inhibitors, such as E-64d, also inhibited coronaviral entry in certain cell types, suggesting a role for cathepsin L and possibly other cysteine proteases (18, 20, 22). Most recently, Riva *et. al* demonstrated in a drug repositioning screen that various cathepsin inhibitors have the ability to block SARS-CoV-2 entry into cells at sub-micromolar concentrations (23).

**K777** (also known as, K11777, S-001, SLV213, 4-methyl-N-((S)-1-oxo-3-phenyl-1-(((S)-1-phenyl-5- (phenylsulfonyl)pentan-3-yl)amino)propan-2-yl)piperazine-1-carboxamide) is a highly potent, irreversible, covalent inhibitor of mammalian cathepsin L and other cysteine proteases of clan CA (Figure 1) (24, 25).

**Figure 1.**
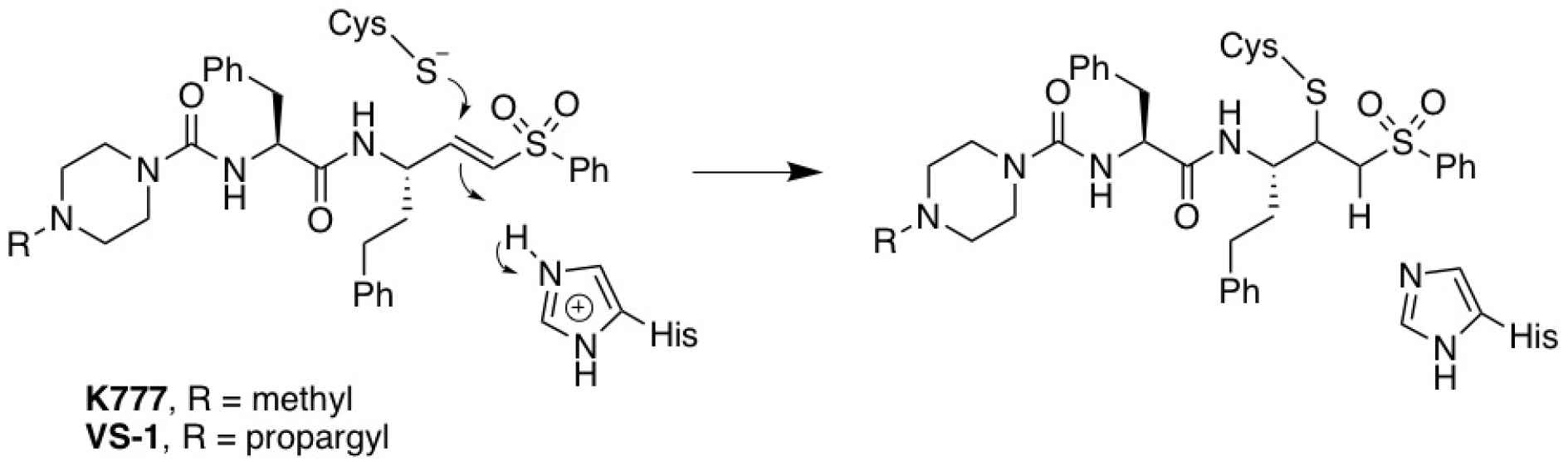
Structure of K777 and its N4-propargyl analogue (VS-1 or K777-alkyne; (26)). Chemical mechanism of irreversible covalent inactivation of Clan CA cysteine proteases.

K777 was originally developed as an inhibitor of cathepsin S,(24) but later showed promise as an anti-parasitic agent (25–31). **K777** has since been approved by the FDA as a clinical stage compound (Selva Therapeutics, Del Mar, CA).

In 2015, Zhou and colleagues reported that **K777** blocked entry of pseudovirus forms of SARS-CoV-1 and MERS into Vero E6 or HEK293 cells, likely due to inactivation of cathepsin L(CTSL) on cell surfaces and/or within endosomes (21). These results suggest that **K777** might also be a potential therapeutic agent for the treatment of COVID-19. Here we show that **K777** blocks CTSL-mediated processing of the spike protein and blocks infection of SARS-CoV-2 in several mammalian cell lines.

## RESULTS

### K777 blocks SARS CoV-2 infectivity of host cells

The anti-coronarial activity of K777 was evaluated in multiple, relevant infected mammalian cell lines (Fig. 2 and Table 1). EC_50_ values for inhibition of viral invasion of host cells by K777 was assessed by observation of a cytopathic effect (CPE). The concentration dependence of the anti-CoV-2 effects of K777 are shown for three cell lines in Fig. 2, in which K777 was most active vs. HeLa cells which express the ACE2 receptor (EC_50_ = 4 nM). For Caco-2 cells virus titer was used as CPE was not observed. EC_50_ values ranged from 4 nM for HeLa/ACE2 cells to 5.5 μM for Calu-3 cells and 4.3 μM for Caco-2 cells. Note that the 2B4 cells, which are Calu-3 cells that have been selected for higher expression of the ACE2 receptor, were more potently inhibited by K777 than standard Calu-3 cells obtained from the ATCC. A549/ACE2 and HeLa/ACE2 cells were the most susceptible to blockage of viral infection by K777. No host cell toxicity was seen for any cell line at inhibitor concentrations of 10-100 μM.

**Table 1.** Anti-CoV-2 Activity in Primate and Human Cells.

| Cell Line | Cell Type | Coronavirus | Laboratory | Inhibition of Viral Infectivity<br>EC<sub>50</sub> (μM) | Cytotoxicity to Host Cells<br>CC<sub>50</sub> (μM) |
| --- | --- | --- | --- | --- | --- |
| Vero E6 | African Green Monkey / kidney epithelial | SARS CoV-1* | UTMB | 2.5 | >10 |
| Vero E6 |  | MERS* | UTMB | 0.62 | >10 |
| Vero E6 |  | SARS CoV-2* | UTMB | 0.62 | >10 |
| Vero E6 |  | SARS CoV-2 | USU | <0.1 | >10 |
| Vero E6 |  | SARS CoV-2 | UCSD | 0.07 | >10 |
| Caco-2 | Human colon carcinoma | SARS CoV-2 | USU | 4.3 | >10 |
| A549/ACE-2 | Human lung carcinoma expressing ACE-2 receptor | SARS CoV-1** | UTMB | 0.08 | >10 |
| A549/ACE-2 |  | MERS** |  | 0.08 | >10 |
| A549/ACE-2 |  | SARS CoV-2** |  | <0.08 | >10 |
| HeLa/ACE-2 | Human cancer cells expressing ACE-2 receptor | SARS CoV-2 | UCSD | 0.004 | >10 |
| Calu-3/2B4 | Human lung epithelial cells | SARS CoV-2 | UTMB | 1 | >10 |
| Calu-3 | Human lung epithelial cells | SARS CoV-2 | UCSD | 3.7 | >10 |

Infected with 100\* or 500\*\* viral particles/sample

**Figure 2.**
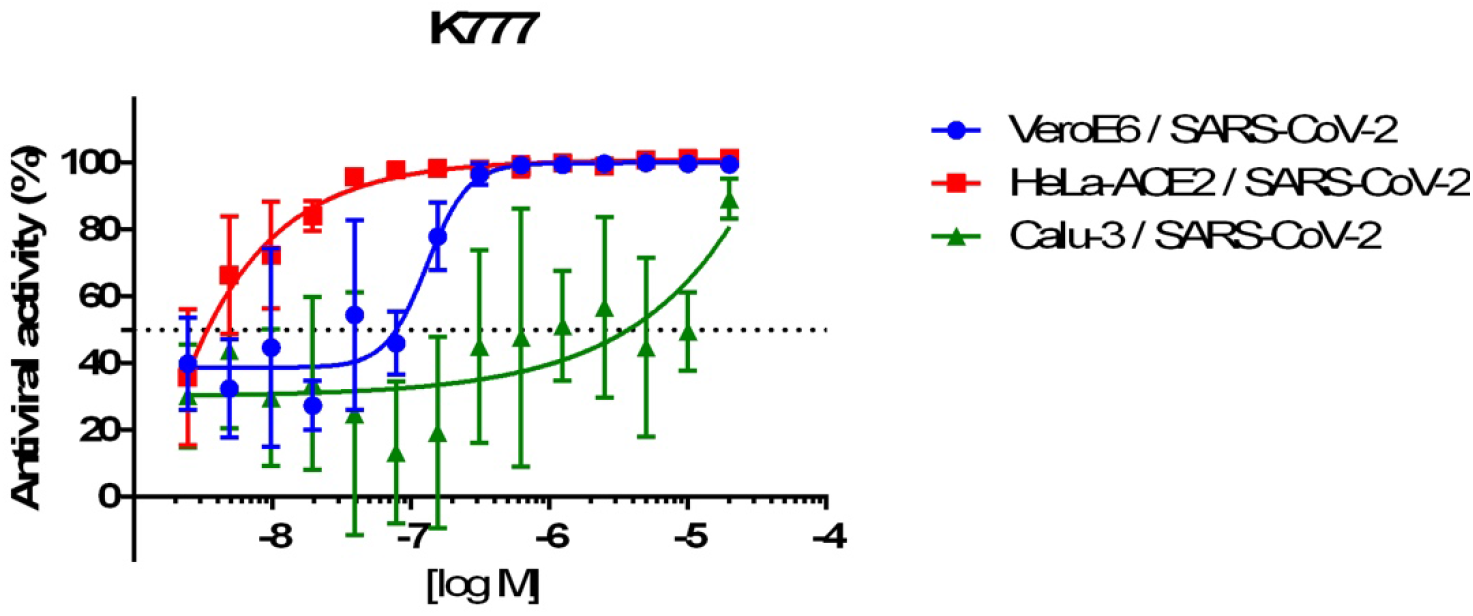
Dose dilution curves for K777 effect on virus infection of three cell lines.

### Analysis of K777 versus SARS-CoV-2 and Clan CA cysteine proteases

In assays carried out at both UC San Diego and Texas A&M University, there was no inhibition of either the PL or the 3CL proteases of SARS-CoV-2 (25-100 nM enzyme concentration) up to 100 μM K777 (Table 2; Figure S5). Conversely, K777 exhibited potent activity against human cathepsin L, where the second-order rate constant of inactivation, *k*_inact_/*k*_I_, was found to be 50-fold and 333-fold greater than cathepsin K and cathepsin B, respectively (Table 2, Figs. S1-S4). K777 formed irreversible covalent adducts with these enzymes (Fig. S3), but exhibited weak, slowly reversible, inhibition of cathepsin C and potent, but rapidly reversible inhibition of cathepsin S (Fig. S4).

**Table 2.** Inactivation or inhibition of mammalian and CoV-2 cysteine proteases^*a*^.

| Enzyme | $K_I$ (nM) | $k_{\text{inact}}$ (s<sup>-1</sup>) | $k_{\text{inact}}/K_I$ (M<sup>-1</sup> s<sup>-1</sup>) |
| --- | --- | --- | --- |
| Human Cathepsin L | 5 ± 2 | 0.013 ± 0.002 | (3 ± 1) × 10<sup>6</sup> |
| Human Cathepsin K | 400 ± 100 | 0.022 ± 0.004 | (6 ± 2) × 10<sup>4</sup> |
| Human Cathepsin B | 3,000 ± 1000 | 0.026 ± 0.007 | (9 ± 4) × 10<sup>3</sup> |
| Bovine Cathepsin C | >100,000 | ND | (1.2 ± 0.4) × 10<sup>1</sup> † |
| Human Cathepsin S | 2.0 ± 0.1 ‡ | ND | ND |
| SARS-CoV-2 3CLpro | >100,000 | ND | ND |
| SARS-CoV-2 PL | >100,000 | ND | ND |

<sup>a</sup> pH = 5.5 for cathepsins and pH 7.0 for CoV-2 proteases; ND, not determinable.
† Value approximated from slope of $k_{\text{obs}}$ vs inhibitor
‡ Value represents overall dissociation constant.

### Activity-Based Protein Profiling of K777 protein targets in Vero E6 cells

We prepared a K777 analog, N4-propargyl-K777 (K777 Alkyne; Fig. 1, (26)) to elucidate the cellular target(s) of K777. This K777 probe facilitates copper-emdiated cycloaddition between an azide and alkyne to both enrich target proteins for mass spectrometry analysis, and for visualization of target proteins via in-gel fluorescence. In both virally and non-virally infected cells, the protein(s) bound to the K777-Cy7 conjugate had a molecular weight of approximately 25 kDa (Fig. 3A), a molecular weight that could correspond to either cathepsin B (mature form: 25 kDa) or cathepsin L (mature form: 24.2 kDa). To confirm that the specificity of the K777 alkyne was analogous to that of K777, recealllsso we preincubated with 1 μM K777 prior to treatment with the K777 alkyne probe. This resulted in >80% blockage of the 25-kDa target protein signal in both virally and non-virally infected cells (Fig 3B). As the other minor protein bands visualized were unchanged upon K777 pre-treatment, it is clear these proteins are not targets of K777.

**Figure 3.**
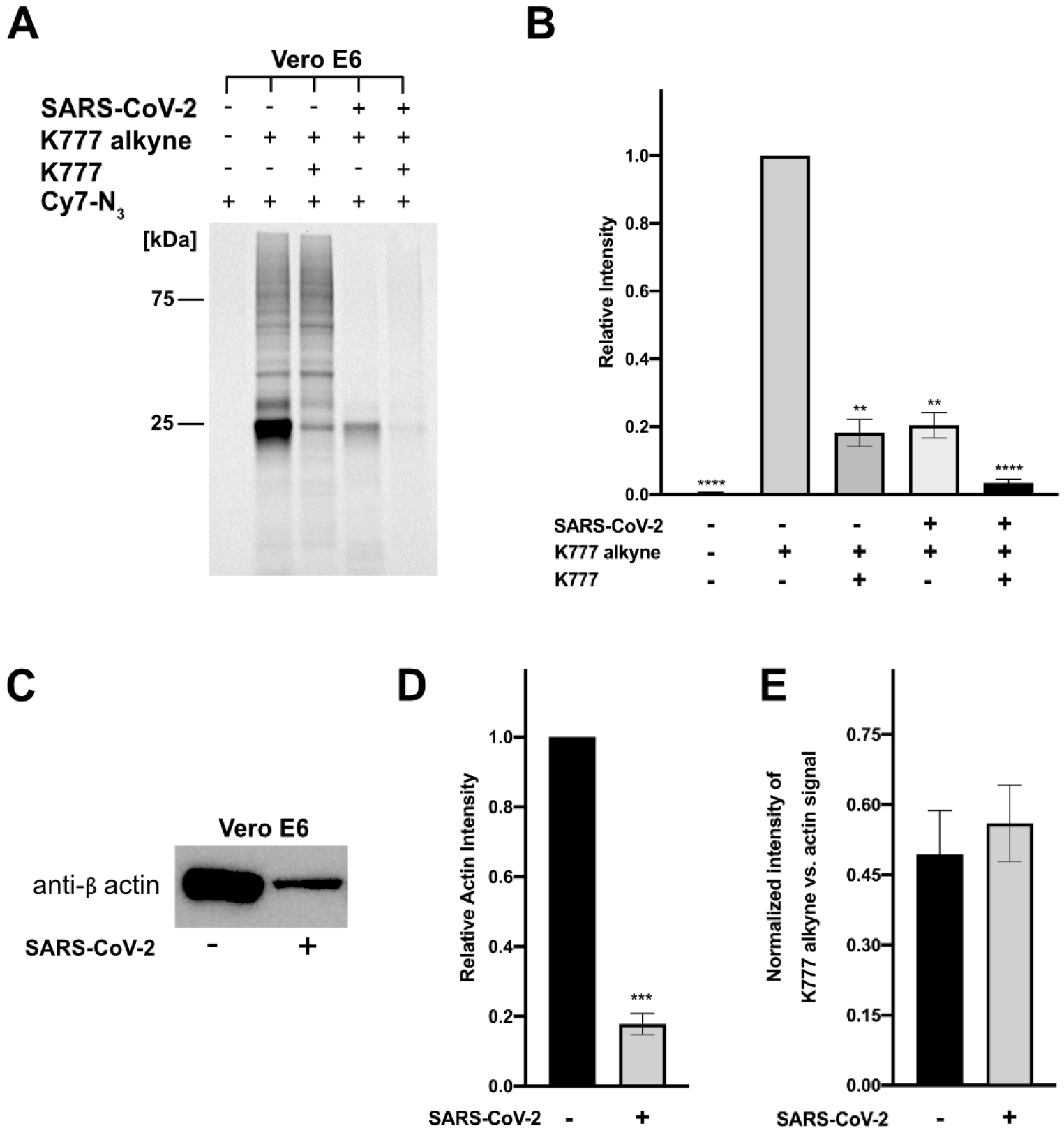
K777 alkyne specifically targets a non-viral protein in SARS-CoV-2 infected and uninfected Vero E6 cells. **A.** Cy7 azide labeled proteins (SARS-CoV-2 infected and uninfected) at 1 μM K777 alkyne is blocked by pre-treatment with 1 μM K777. **B.** Densitometry analysis of **A,** and a replicate in gel fluorescence experiment. **C**. Blotting of total β-actin levels in SARS-CoV-2 infected and non-infected cells. **D.** Densitometry analysis of relative actin levels. **E.** Comparison of the signal intensity of the 25 kDa band enriched in the presence of K777 alkyne (SARS-CoV-2 infected and uninfected) versus the β-actin signal (SARS-CoV-2 infected and unifected).

When comparing the intensity of the 25-kDa band in SARS-CoV-2-infected cells to that of its uninfected counterpart there was nearly a 5-fold decrease in intensity (Figure 3A, 3B). Reduction of the signal for this band in the virally-infected samples may be due to either a reduction of total protein or differential expression of the target(s) of the K777-alkyne. To investigate this, we immunoblotted the housekeeping protein human β-actin to compare its relative amounts in both sets of samples. Upon quantifying the difference in β-actin content between the SARS-CoV-2-infected and uninfected cells, we measured a 5.7-fold decrease in total β-actin signal in the Vero E6 SARS-CoV-2 infected samples (Figure 3C, 3D; S6, S7). Comparison of the relative intensities of the β-actin band versus the 25-kDa enriched protein for both SARS-CoV-2 infected and uninfected cells approximated a 1:1 ratio (Figure 3E). These results suggested that coronaviral infection of Vero E6 caused a global reduction in protein expression, in agreement with recent reports that NSP1 of SARS-CoV-2 suppresses protein translation in infected cells(34).

### Identification of Cathepsin B and L as K777 targets by affinity enrichment and proteomic analysis

To identify the molecular target(s) of the K777 alkyne, affinity enrichment and proteomic analysis of treated Vero E6 cell lysates was carried out with variable concentrations of the K777 alkyne (0-10 μM) (Fig. S8A). 117 proteins at a false discovery rate of 1% were identified, of which 65 were present in the K777 alkyne-treated samples, but not the untreated control (Fig. S8B). Proteins of the highest abundance in the treated (5 and 10 μM samples), as determined by MaxQuant LFQ analysis (35), were cathepsin B (CTSB) and cathepsin L (CTSL).

### Cathepsin L but not cathepsin B catalyzes cleavage of the SARS CoV-2 spike protein

Having identified CTSB and CTSL as putative targets of K777 in cells, we sought to elucidate the ability of these proteins to process the SARS-CoV-1 and SARS-CoV-2 spike proteins *in vitro*. We determined that both proteases could process the SARS-CoV-1 spike protein, but only CTSL could process the SARS-CoV-2 spike protein (Figure 4A, 4B, S9). Additionally, processing of SARS-CoV-2 spike protein by CSTL produced different sized molecular weight bands than that of SARS-CoV-1 spike protein. Immunoblotting against the C-terminal Flag epitope of the SARS-CoV-2 S resulted in immuno-reactivity with both the intact S protein and a ~120 kDa protein fragment, indicating that this ~120 kDa protein contained an intact S2 domain, and that CTSL proteolysis took place in the S1 domain of the spike protein at an unknown site, which we have coined S1’ (Figure 4C). K777 efficently blocked the cleavage at this S1’ site by CTSL of SARS-CoV-2 (Figure 4D).

**Figure 4.**
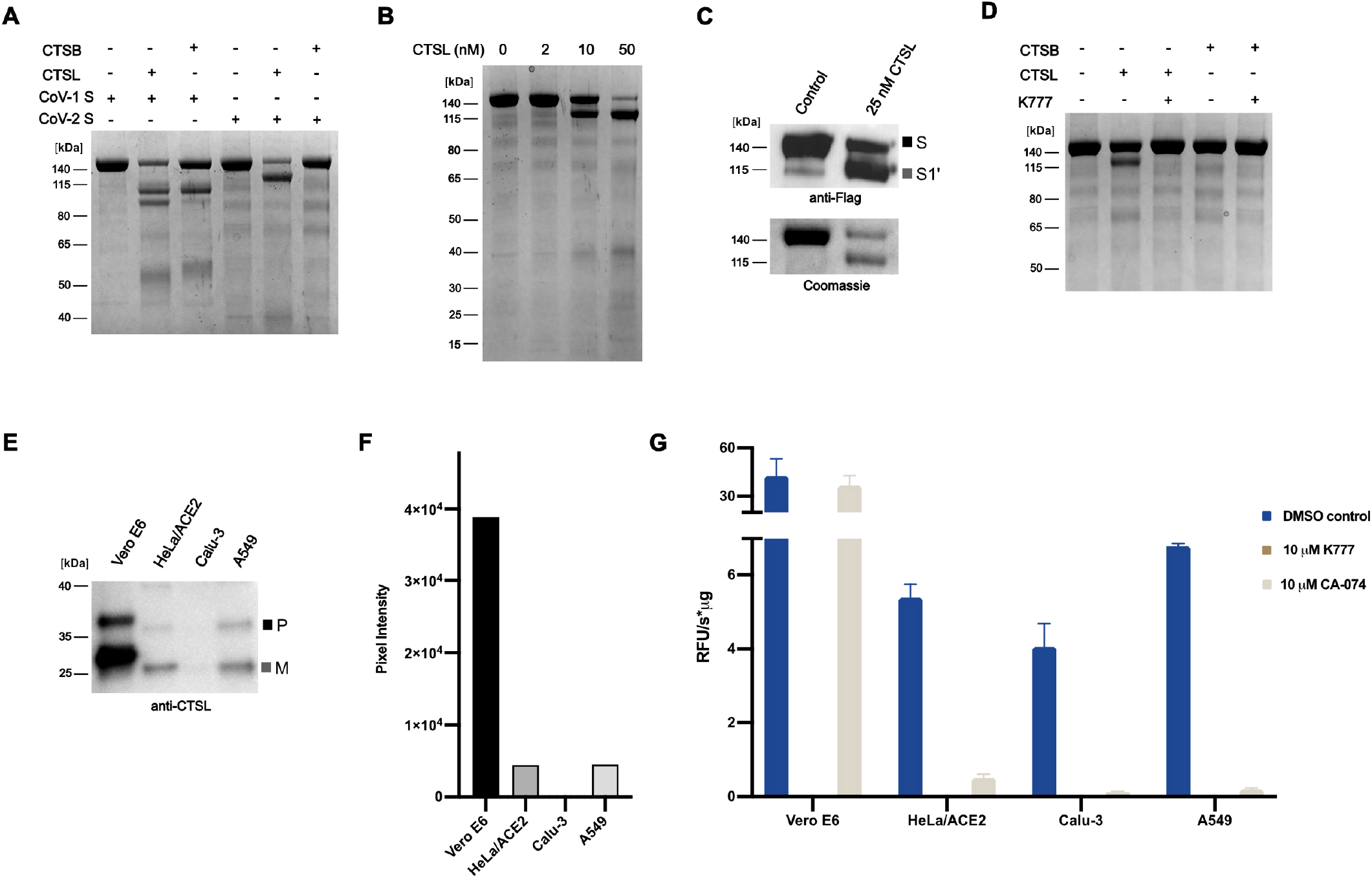
CTSL and CTSB catalyze differential processing of the SARS-CoV-2 spike protein and CTSL levels vary in cells. **A**. Processing of SARS-CoV-1 spike protein and SARS-CoV-2 spike protein by CTSB (250 nM) and CTSL (25 nM) **B**. Concentration-dependent cleavage of the SARS-CoV-2 spike protein by CTSL. **C**. Western blotting against the C-terminal FLAG epitope of SARS-CoV-2 spike demonstrates that CTSL cleavage occurs in the S1 domain. **D.** CTSL (25 nM) proteolysis of the SARS-CoV-2 spike protein is inhibited by K777. **E**. Western blotting against CTSL shows differential expression between cell lines (P: pro form of CTSL; M: mature form of CTSL). **F.** Densitometry analysis of **E**. **G**. Activity of cell lysate with Z-FR-AMC in the presence and absence of inhibitors.

### Cathepsin L is present in all cell lines tested but levels vary

Although K777 targets both CTSL and CTSB, inhibition of CTSL is apparently responsible for its potent anti-viral effects. To investigate why virally-infected A549/ACE2, HeLa/ACE2, and Vero E6 cells are more sensitive to K777 treatment than Calu-3 cells, we evaluated the relative abundance of CTSL protein in each cell line. Human CTSL shares 96% protein sequence identity with CTSL in African green monkeys (Fig. S10) and therefore both the proprotein (upper band) and mature active enzyme (lower band) can be detected by immunoblot using an anti-human CTSL antibody (Figure 4E-F). Mature CTSL was 8.8-fold and 8.7-fold higher in Vero E6 extracts compared to HeLa/ACE2 and A549/ACE2, respectively. Under these exposure conditions, CTSL was not detectable in Calu-3 cell extracts. The large difference in CTSL abundance between these cell lines was further validated by quantitative proteomic analysis using a CTSL-derived peptide (NHCGIASSASYPTV) that is identical in humans and African green monkeys (Fig. S10). This peptide was between 32- and 57-fold more abundant in Vero E6 compared to the other three cell lines, therefore supporting the immunoblotting studies. Finally, we quantified the specific activity of CTSL in all four cell lines using the fluorogenic substrate Z-FR-AMC. The cell extracts were first treated with broad spectrum serine, aspartic acid and metallo-protease inhibitors to inactivate non-cysteine proteases that may hydrolyze this substrate. The remaining activity in each cell line was eliminated by K777 confirming that only cysteine proteases were active in the non-K777 treated samples. As K777 inhibits both CTSL and CTSB, we treated the cell lines with CA-074, a selective CTSB inhibitor. Cysteine protease activity in A549/ACE2, HeLa/ACE2 and Calu-3 cell lystates was found to be mostly due to CTSB, while activity in Vero E6 cells was mostly CTSL. The specific activity of CTSL was determined to be 0.1, 0.2, 0.5 and 36.7 RFU s^−1^ μg^−1^ for Calu-3, A549/ACE2, HeLa/ACE2, and Vero E6, respectively (Figure 4G). These studies showed that the concentration of CTSL in HeLa/ACE2, A549/ACE2 and Vero E6 cells strongly correlated with the EC_50_ values generated for K777 treatment of SARS-CoV-2 infected cells. However, Calu-3 cells had the lowest levels of CTSL but the highest EC_50_ values, suggesting that another protease may be involved in spike protein processing.

## DISCUSSION

Three host proteases, TMPRSS2, furin, and CTSL, have been proposed as key enzymes for the processing of the coronaviral spike protein (18, 20–22). In our analysis, we have shown that K777, a potent inhibitor of CTSL activity, is also a potent anti-SARS-CoV-2 infection agent in several host cell models. Although the maximal rates of inactivation, *k*_inact_, for cathepsins L, B, and K are similar, the second-order rate constant of time-dependent inactivation (*k*_*inact*_/K_I_) varies by over two orders-of-magnitude for these enzymes and by five orders-of-magnitude compared to cathepsin C, highlighting the functional selectivity of K777 for CTSL versus other host-cell cathepsins. Using a propargyl analog of K777 as an affinity probe we determined that CTSB and CTSL were intracellular targets of this compound in Vero E6 cells, however only CSTL, but not CTSB, cleaved the SARS-CoV-2 spike protein. This cleavage occurred in the S1 domain of the spike protein at a site different from that observed in the S protein of SARS-CoV-1. The S1 domains of the spike proteins from SARS-CoV-1 and SARS-CoV-2 share 64% sequence identity, and analysis of the SARS-CoV-2 spike protein structure revealed multiple unresolved loop regions in the S1 domain, suggesting that CTSL proteolysis may occur in one of these unstructured regions (36, 37)

K777 inhibited viral entry into three host cell models including cell lines derived from African green monkey renal epithelium (Vero E6), human cervical epithelium (HeLa/ACE2) and human lung epithelium (A549/ACE2) with nanomolar potency. Anti-viral inhibition was lower (EC_50_ = 3.7 and 4.3 μM) in a second human lung epithelium cell line (Calu-3) and a human colon epithelium-derived cell line (Caco-2). Why is the efficacy of K777 different in different cell lines? We quantified differential levels of CTSL in cell lines via Western blotting and an enzyme activity assay. Vero E6 cells expressed the highest levels of CTSL and HeLa/ACE2 and A549/ACE2 had lower amounts. These data suggest that the amount of CTSL correlates with K777 efficacy. However, Calu-3 cells have the lowest amount of CTSL and yet are relatively insensitive to K777 treatment. Previous work has shown that treatment of MERS-infected Calu-3 cells with E64d, a broad cysteine protease inhibitor, had diminished efficacy presumably as a result of low CTSL expression but heightened TMPRSS2 expression. This shift in expression profile resulted in the predominant processing of the MERS spike protein by TMPRSS2 (38). It was also suggested that cleavage of the SARS-CoV-2 spike protein by TMPRSS2 in cells with low CTSL levels (such as Calu-3) is essential for efficient viral entry, whereas in TMPRSS2 negative cells (Vero E6) viral infection can proceed with high efficiency using CTSL (39). On the other hand, Hoffman and co-workers(18) concluded that the TMPRSS2 inhibitor, camostat mesylate, reduced but did non abrogate SARS-CoV-2 infection of Calu-3 cells. They suggested that following camostat treatment, there remains residual spike protein that might require processing by CTSL. The EC_50_ for camostat mesylate inhibition of SARS-CoV-2 infection of Calu-3 cells was 1 uM, in the same range as K777(Table 1). Therefore, future experiments should evaluate a combination of K777 and an appropriate TMPRSS2 or furin inhibitor. Combination therapy might protect all host cells from viral infection even if some utilize both CTSL and TMPRSS2 (or furin) for processing of the SARS-CoV-2 spike protein.

In summary, there are currently two potential therapeutic uses of K777. First, it may block primary infection by SARS-CoV-2. If primary infection occurs in lung epithelium, it could be argued that the reduced activity of K777 to prevent infection of Calu-3 cells is concerning. However it should be noted that the anti-viral infection activity of K777 was robust in another lung epithelial cell line, A549/ACE2 (Table 1). Furthermore both Calu-3 and A549/ACE2 cells are derived from adenocarcinomas of the lung, not primary lung epithelium. Adenocarcinomas have multiple mutations compared to their cell of origin and that may alter infection potential. Alternatively, K777 might be employed to prevent viral infection of other host organs if spike protein processing in cells of those organs depends on CTSL. Confirmation of these suggestions will come from animal model testing and ultimately human clinical trials. Support for the rapid advancement of K777 as potential COVID-19 therapy comes from the fact that K777 is orally bioavailable, and safety, tolerability and PK/PD have been evaluated in several preclinical model systems, including non-human primates (31). An IND application for K777 was recently approved by the FDA(Selva Therapeutics, Del Mar, CA).

## MATERIALS AND METHODS

### Chemicals and Proteins

The synthesis of K777, its N-propargyl analog (25), the 3CLpro FRET substrate, and information regarding the expression, and purification of 3CLpro (Mainpro) are provided in the Supplementary Appendix. Cyanine7-azide was obtained from Click Chemistry Tools. Recombinant proteases were obtained from the following vendors: human cathepsin L (Millipore Sigma, Athens Research and Technology, Inc., (Texas A&M) or R&D Systems (UCSD), SARS-CoV-2 PLpro (Acro Biosystems, PAE-C5184), human cathepsin S (Millipore Sigma), bovine spleen cathepsin C (Millipore Sigma), human liver cathepsin B (Millipore Sigma), and human cathepsin K (Enzo Sciences, Inc.). Substrates were purchased from the following vendors: Z-FR-AMC (EMD Millipore), GF-AMC (MP Biomedicals), and Z-LR-AMC (Enzo Life Sciences, Inc.). Recombinant SARS-CoV-1 spike protein was obtained from SinoBiological and SARS-CoV-2 spike was obtained from Genscript and Acro Biosystems.

### SARS-CoV-2 protease assays

Recombinant SARS-CoV-2 3CLpro (100 nM) was assayed at 25°C in either (a) in 30 μL reaction volumes containing 20 mM Tris-HCl pH 7.5, 150 mM NaCl, 1 mM DTT, 5 % glycerol, 0.01% Tween 20 100 μM of Mu-HSSKLQ-AMC (Sigma-Aldrich, SCP0224), in 384-well black plates in triplicate or (b) in 50 μL reaction mixtures containing 20 mM Tris-HCl pH 7.5, 150 mM NaCl, 0.1 mM EDTA, 2 mM DTT, 10% DMSO, and 25-50 nM 3CLpro in 96-well plates (Greiner, flat-bottom half volume, clear black plates), using the FRET-based substrate Abz-SAVLQSGFRK(DNP)-NH_2_ wherein peptidolysis was measured at 320/420 nm (ex/em) (Biotek® Synergy M2) in the presence and absence of 0-50 μM K777 in duplicate. Recombinant SARS-CoV-2 PL was assayed in either (a) 50 mM HEPES, 150 mM NaCl, 1 mM DTT, 0.01% Tween 20; pH 6.5 buffer using 50 μM Z-RLRGG-AMC (Bachem, I1690) and 24.5 nM enzyme or (b) 50 mM HEPES pH 7.5, 5 mM DTT, 0.1 mg/mL BSA, 2% DMSO, 50 μM Z-RLRGG-AMC, and 10 nM PL, where rates in the presence and absence of K777 were measured at 360/460 nm (ex/em). In some studies, enzymes were pre-incubated with 20 μM of K777 or 0.0025 % DMSO for 15 minutes and then diluted in an equal volume of substrate in assay buffer.

### Analysis of K777 as an inhibitor of Clan CA cysteine proteases

Assays were conducted at 25°C in 50 mM sodium acetate (pH 5.5), 1 mM EDTA, 1 mM CHAPS, 10% (v/v) DMSO, and 5 mM DTT containing 1 nM cathepsin B, 30-50 pM cathepsin S, 320 pM cathepsin K, 1 nM cathepsin L, and 132 nM cathepsin C in the presence of 0-0.1 mM K777 and 0.01 mM Z-FR-AMC for cathepsin S, cathepsin B, and cathepsin L, GF-AMC for cathepsin C, and Z-LR-AMC for cathepsin K) 96-well black microplate (Corning Costar®). Rates of peptidolysis of the dipeptide-AMC substrate(s) were measured at either a SpectraMax M5 (Molecular Devices) in or a Synergy HTX (Biotek, Wisnooki, VT) microplate reader with λ_ex_ = 360 nm, λ_em_ = 460 nm in 8-60 sec intervals. The initial velocity in relative fluorescent units per second was calculated for 10 minutes.

To determine the mechanism of inhibition, 10 nM cathepsin L or 2.5 nM cathepsin S was incubated with 10 nM K777, 17.5 nM cathepsin B or 3.75 nM cathepsin K with 200 nM K777, and 1.49 μM cathepsin C with 100 μM K777 for 1 h at room temperature, followed by 50-fold dilution into assay buffer and their residual activity was measured under nearly identical conditions to assay described above.

### Cell cultures and SARS-CoV-2 infection

#### University of Texas Medical Branch

African green monkey derived cells Vero E6 [CRL:1586, ATCC] were grown in Eagle’s minimal essential medium (EMEM) supplemented with standard doses of penicillin and streptomycin, and 10% fetal bovine serum (FBS), designated M-10 medium. Human A549 cells stably transduced with human ACE2 viral receptor (A549/ACE2) and Calu-3 (2B4 clones selected for increased ACE2 receptor) cells were grown in M-10 and M-20 medium, respectively. SARS-CoV-2 (USA_WA1/2020 isolate), the 3rd passage in Vero E6 cells from the original CDC (Atlanta) material and sequence confirmed, was used throughout the study. All experiments using infectious virus were conducted at the University of Texas Medical Branch under BSL-3 conditions.

#### UC San Diego

Vero E6 cells (CRL-1586, ATCC) were cultivated in DMEM in a similar manner as that described above. Calu-3 cells (HTB-55, ATCC) were cultivated in MEM media supplemented with 10% FBS, penicillin (100 units/ml) and streptomycin (100 μg/ml) and 1x GlutaMAX (Fisher Scientific). The SARS-CoV-2 virus used was the USA-WA1/2020 isolate.

#### Utah State University

Vero E6 cells [CRL:1587, ATCC] or Caco-2 cells [HTB-37, ATCC] were grown in minimum essential medium (MEM) containing 5% FBS or 10% FBS respectively. The same SARS-CoV-2 (USA_WA1/2020 isolate) was used.

### Evaluation of protease inhibitors in a coronaviral infection assay

#### University of Texas Medical Branch

A modified Vero E6-based standard micro-neutralization assay was used to rapidly evaluate the drug efficacy against SARS-CoV-2 infection. Briefly, confluent Vero E6 cells grown in 96-wells microtiter plates were pre-treated with 8 nM to 25 μM K177 (2-fold serially diluted) for 2 h before infection with ~100 infectious SARS-CoV-2 particles in 100 μl EMEM supplemented with 2% FBS. Vero E6 cells treated with 2-fold serially diluted dimethyl sulfoxide (DMSO) with or without virus were included as positive and negative controls, respectively. A549/ACE2 and Calu-3/2B4 cells were treated with 500 and 100 viral particles, respectively, prior to addition of K777. After cultivation at 37°C for 4 days, individual wells were observed under the microcopy for the status of virus-induced formation of cytopathic effect (CPE). The efficacy of individual drugs was calculated and expressed as the lowest concentration capable of completely preventing virus-induced CPE in 100% of the wells. All compounds were dissolved in 100% DMSO as 10 mM stock solutions and diluted in culture media.

#### UC San Diego

For Vero E6 and HeLa/ACE2, 1,000 cells were seeded per well, and for Calu-3, 3,000 cells/well. The plates were incubated for 2 h in the incubator at 37°C, 5% CO_2_. SARS-CoV-2 was added to the plate wells in a multiplicity of infection (MoI) of 1 in a final volume of 25 μL. The plates were again incubated for 24 h at 37°C and 5% CO_2_, followed by the addition of 25 μl/well of 8% paraformaldehyde solution. Cells were fixed by incubation at room temperature for 30 min. The content was then aspirated and the wells incubated with 1:1,000 dilution of the Rabbit IgG antibody against SARS-CoV-2 capsid (Genetex, GTX135357) for 1 h and washed. Rabbitt IgG coupled with AlexaFluor488 diluted 1:200 was added to the wells and incubated for at least 2 h. To calculate the antiviral activity, the average infection ratio from the untreated controls (0.1% DMSO) were normalized as 0% antiviral activity. The average infection ratio from the uninfected controls (no SARS-CoV-2) were normalized to 100% antiviral activity. A linear regression was applied to calculate the antiviral activity of each well related to the normalized controls. Serial dilution tests were performed to assess the antiviral effect and potency (IC_50_) of K777 in the three cell lines.

#### Utah State University

SARS-CoV-2 CPE and virus yield reduction (VYR) assay were carried out as follows: confluent cell culture monolayers of Vero E6 or Caco-2 cells were prepared in a 96-well disposable microplate the day before testing. An insufficient CPE was observed on Caco-2 cells to use for calculation of the EC_50_ of the CPE so data were obtained after 2 hours of treatment at 37ºC. CPE values were normalized based upon virus controls and 50% effective (EC_50_) and 50% cytotoxic (CC_50_) concentrations were determined using non-linear regression.

### Identification of protein targets of K777 using an activity-based probe, 4N-propargyl-K777

4N-propargyl-K777 was added at concentrations of 1 μM to Vero E6 cells, with and without infection by SARS CoV-2 viral particles at an MoI of 0.1. For samples treated with both K777 and the K777 alkyne, Vero E6 cells were first pre-treated with K777 (1 μM) for 1 h, prior to the addition of 4N-propargyl-K777 (1 μM). Cell growth continued for 4 days, followed by harvest, and ~ 20 million cells were suspended in 1 mL of lysis buffer (200 mM Tris, 4% CHAPS, 1M NaCl, 8 M Urea, pH 8.0) and protease inhibitor cocktail (Sigma Aldrich, #8340 at a ratio of 50:1 (v/v). Cells were lysed by sonication (ThermoFisher), protein concentration quantified by BCA (Pierce), followed by copper-mediated click-chemistry with a Cy7-azide (Click Chemistry Tools) as described in the SI appendix, followed by SDS-PAGE separation (Invitrogen) and imaging (ChemiDoc imager, Bio-Rad).

### Azide Enrichment and Proteomic analysis of Vero E6 K777-alkyne treated cells

4N-propargyl-K777 (alkyne K777, (26)) was added at concentrations of 0-10 μM to Vero E6. Cell lysates were prepared as described above. Enrichment of alkyne-modified proteins was conducted using the Click-&-Go Protein Enrichment Kit (Click Chemistry Tools) and the protocols thereof. The resin bound proteins were reduced with DTT, alkylated with iodoacetamide, and digested on resin using 1 μg of Trypsin Gold (Promega) at 37°C overnight with agitation. The digested peptides were eluted from the column, desalted (Pierce™ C18 Tips, Thermo Fisher Scientific), concentrated to dryness, and suspended in 25 mM ammonium bicarbonate followed by nanoLC-MS/MS analysis on an Orbitrap Fusion™ Tribrid™ Mass Spectrometer (ThermoFisher). For more details refer to the SI Appendix.

### Western Blotting

#### Texas A&M University

Cell lysates were prepared as stated above. Samples were separated via SDS-PAGE (Invitrogen) and semi-dry transferred (Bio-Rad) onto an activated PDVF membrane (GE Healthcare Life Sciences) followed by blocking with TBST/5% wt/vol non-fat milk (1 hr, RT). Anti-actin antibody (Sigma Aldrich) or anti-Flag (Cell Signaling Technologies) was added at a 1:100,000 or 1:25,000 dilution respectively and incubated overnight at 4°C. The membrane was washed 5x with TBST, probed with goat-anti rabbit HRP conjugate antibody (1:3000) (Bio-Rad) in TBST/5% wt/vol non-fat milk (1 h, RT), washed (5x, TBST) followed by addition of ECL substrate (Bio-Rad) and imaged (ChemiDoc, Bio-Rad).

#### UC San Diego

Protein extracts from Vero-E6, HeLa/ACE2, Calu-3 and A549/ACE2 cell lines were prepared by sonication in 50 mM MES, pH 6.0, containing 1 mM EDTA, 1 μM pepstatin, 100 μM AEBSF and 1 mM DTT in an ice bath. The extracts were cleared by centrifugation (16,000 g at 4°C for 10 min) and the protein concentration was determined by BCA assay (Pierce). 10 μg of protein were resolved by SDS-PAGE (15% polyacrylamide gel) under reducing conditions, transferred to a PVDF membrane and blocked for 16 h in 10% non-fat milk in TBST. The membrane was then washed 3x in TBST and incubated for 1 h with anti-CatL polyclonal IgG (R&D Systems, AF952-SP) diluted 1:1,000 in TBST followed by 30 min with rabbit anti-goat HRP IgG (SouthernBiotech, #6164-05) at a dilution of 1:2,000. After washing in TBST, the membrane was developed with Immobilon Crescendo Western HRP substrate (MilliporeSigma) and imaged using an ChemiDoc Imaging System (Biorad).

### Mammalian protease activity assays

2.5 μg of SARS-CoV-1 and SARS-CoV-2 spike protein was incubated with trypsin (25 nM) in 50 mM HEPES, 1 mM Na-EDTA, pH 7.5 or with cathepsin B (250 nM), or cathepsin L (2-250 nM) in 50 mM sodium acetate, 1 mM Na-EDTA, 1 mM CHAPS, and 1 mM DTT, pH 5.5. Inhibitors were solubilized in DMSO and added to 10% v/v DMSO (2.5-5 μM K777 and 100 μM leupeptin). After 1 h, reactions were quenched by addition of 4x loading dye containing 5 mM DTT. Samples were denatured at 95°C for 10 min and analysis by SDS-PAGE (Invitrogen). The gels were stained with either Coomassie Blue or transferred to a PDVF membrane for Western blotting as described above, and imaged (ChemiDoc, Bio-Rad). Protein extracts from Vero E6, HeLa/ACE2, Calu-3 and A549/ACE2 cell lysates (225 ng) were incubated for 30 min with 10 μM of K777 or CA-074 (Cayman #24679) in 50 mM Sodium acetate, pH 5.5 containing 1 mM EDTA, 1 μM pepstatin, 100 μM AEBSF and 5 mM DTT. These samples were then assayed with 25 μM Z-FR-AMC at 25°C in triplicate wells of black, flat-bottomed 384-well microplates in a total volume of 30 μL. Fluorescent activity was quantified as outlined above.

### Statistical analysis

All densitometry and gel quantification analysis was conducted using ImageJ(33) and the calculated pixel intensities were normalized accordingly. For analysis of the statistical significance of the changes in pixel intensity we used an unpaired parametric t-test. P values ≤0.05 (*), ≤0.01 (**), ≤0.001 (***)

## Supporting information

Cover Letter

## ACKNOWLEDGEMENTS

Funding for experiments completed at Utah State University was provided by the Respiratory Diseases Branch, National Institute for Allergy and Infectious Diseases, NIH USA (Contract N01-AI-30048). Funding for work in the Meek lab at Texas A&M was from AgriLife Research, Texas A&M University. For the Tseng lab, NIAID, Grant #: R24 AI120942 NPARS-S01. This research was supported by a Career Award for Medical Scientists from the Burroughs Wellcome Fund to A.F.C. The following reagent was deposited by the Centers for Disease Control and Prevention and obtained through BEI Resources, NIAID, NIH: SARS-Related Coronavirus 2, Isolate USA-WA1/2020, NR-52281. Part of the funding for studies performed at UTMB was provided by Selva Therapeutics, Inc.

---

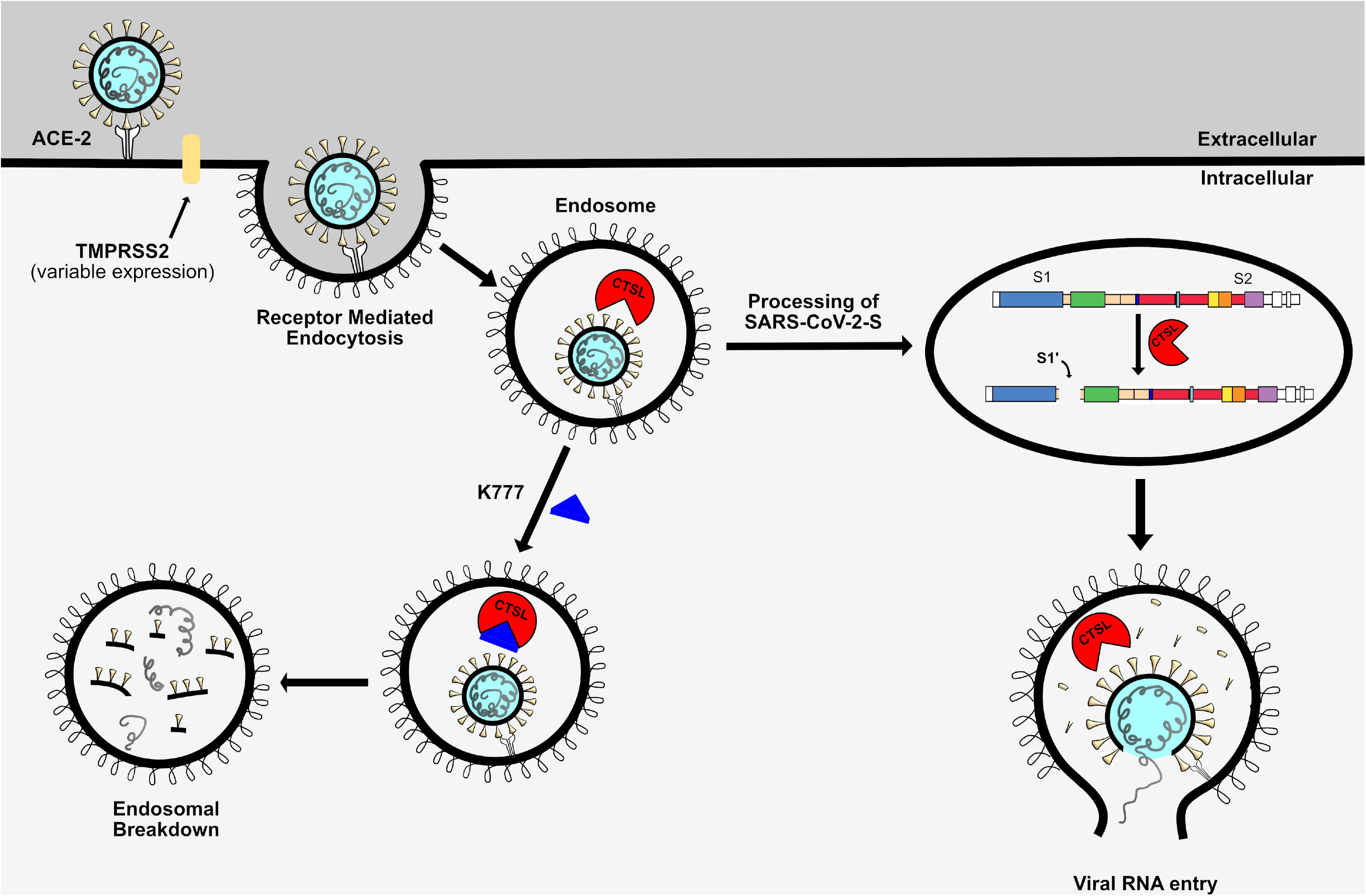

## REFERENCES

1. Zhou P, et al. (2020) A pneumonia outbreak associated with a new coronavirus of probable bat origin. Nature 579(7798):270–273.

2. Morse JS, Lalonde T, Xu S, & Liu WR (2020) Learning from the Past: Possible Urgent Prevention and Treatment Options for Severe Acute Respiratory Infections Caused by 2019-nCoV. ChemBioChem 21(5):730–738.

3. Dong E, Du H, & Gardner L (2020) An interactive web-based dashboard to track COVID-19 in real time. The Lancet Infectious Diseases 20(5):533–534.

4. New York Times (2020) Coronavirus in the U.S.: Latest Map and Case Count.

5. Pillaiyar T, Meenakshisundaram S, & Manickam M (2020) Recent discovery and development of inhibitors targeting coronaviruses. Drug Discovery Today 25(4):668–688.

6. Guy RK, DiPaola RS, Romanelli F, & Dutch RE (2020) Rapid repurposing of drugs for COVID-19. Science 368(6493):829.

7. Rota PA, et al. (2003) Characterization of a Novel Coronavirus Associated with Severe Acute Respiratory Syndrome. Science 300(5624):1394.

8. Zaki AM, van Boheemen S, Bestebroer TM, Osterhaus ADME, & Fouchier RAM (2012) Isolation of a Novel Coronavirus from a Man with Pneumonia in Saudi Arabia. New England Journal of Medicine 367(19):1814–1820.

9. Lu R, et al. (2020) Genomic characterisation and epidemiology of 2019 novel coronavirus: implications for virus origins and receptor binding. Lancet 395(10224):565–574.

10. Zhang L, et al. (2020) Crystal structure of SARS-CoV-2 main protease provides a basis for design of improved α-ketoamide inhibitors. Science 368(6489):409.

11. Báez-Santos YM, St John SE, & Mesecar AD (2015) The SARS-coronavirus papain-like protease: structure, function and inhibition by designed antiviral compounds. Antiviral research 115:21–38.

12. Li L, Li G, Chen J, Liang X, & Li Y (2020) Comparative genomic analysis revealed specific mutation pattern between human coronavirus SARS-CoV-2 and Bat-SARSr-CoV RaTG13. bioRxiv:2020.2002.2027.969006.

13. Heald-Sargent T & Gallagher T (2012) Ready, set, fuse! The coronavirus spike protein and acquisition of fusion competence. Viruses 4(4):557–580.

14. Shang J, et al. (2020) Cell entry mechanisms of SARS-CoV-2. Proceedings of the National Academy of Sciences 117(21):11727.

15. Ou X, et al. (2020) Characterization of spike glycoprotein of SARS-CoV-2 on virus entry and its immune cross-reactivity with SARS-CoV. Nature Communications 11(1):1620.

16. Belouzard S, Chu VC, & Whittaker GR (2009) Activation of the SARS coronavirus spike protein via sequential proteolytic cleavage at two distinct sites. Proceedings of the National Academy of Sciences 106(14):5871.

17. Millet JK & Whittaker GR (2014) Host cell entry of Middle East respiratory syndrome coronavirus after two-step, furin-mediated activation of the spike protein. Proceedings of the National Academy of Sciences 111(42):15214.

18. Hoffmann M, et al. (2020) SARS-CoV-2 Cell Entry Depends on ACE2 and TMPRSS2 and Is Blocked by a Clinically Proven Protease Inhibitor. Cell 181(2):271–280.e278.

19. Huang Y, Yang C, Xu X-f, Xu W, & Liu S-w (2020) Structural and functional properties of SARS-CoV-2 spike protein: potential antivirus drug development for COVID-19. Acta Pharmacologica Sinica 41(9):1141–1149

20. Simmons G, et al. (2005) Inhibitors of cathepsin L prevent severe acute respiratory syndrome coronavirus entry. Proceedings of the National Academy of Sciences of the United States of America 102(33):11876.

21. Zhou Y, et al. (2015) Protease inhibitors targeting coronavirus and filovirus entry. Antiviral Research 116:76–84.

22. Bosch BJ, Bartelink W, & Rottier PJM (2008) Cathepsin L Functionally Cleaves the Severe Acute Respiratory Syndrome Coronavirus Class I Fusion Protein Upstream of Rather than Adjacent to the Fusion Peptide. Journal of Virology 82(17):8887.

23. Riva L, et al (2020) Discovery of SARS-CoV-2 antiviral drugs through large-scale compound repurposing. Nature 586(7827):113–119.

24. Palmer JT, Rasnick D, Klaus JL, & Bromme D (1995) Vinyl Sulfones as Mechanism-Based Cysteine Protease Inhibitors. Journal of Medicinal Chemistry 38(17):3193–3196.

25. Engel JC, Doyle PS, Hsieh I, & McKerrow JH (1998) Cysteine protease inhibitors cure an experimental Trypanosoma cruzi infection. J Exp Med 188(4):725–734.

26. Yang PY, Wang M, He CY, Yao SQ. (2012) Proteomic profiling and potential cellular target identification of K11777, a clinical cysteine protease inhibitor, in *Trypanosoma brucei*. Chem Commun 48(6):835–837.

27. Doyle PS, Zhou YM, Engel JC, & McKerrow JH (2007) A Cysteine Protease Inhibitor Cures Chagas Disease in an Immunodeficient-Mouse Model of Infection. Antimicrobial Agents and Chemotherapy 51(11):3932.

28 Barr SC, et al. (2005) A cysteine protease inhibitor protects dogs from cardiac damage during infection by Trypanosoma cruzi. Antimicrobial agents and chemotherapy 49(12):5160–5161.

29. Abdulla M-H, Lim K-C, Sajid M, McKerrow JH, & Caffrey CR (2007) Schistosomiasis Mansoni: Novel Chemotherapy Using a Cysteine Protease Inhibitor. PLOS Medicine 4(1):e14.

30. Ndao M, et al. (2013) A Cysteine Protease Inhibitor Rescues Mice from a Lethal Cryptosporidium parvum Infection. Antimicrobial Agents and Chemotherapy 57(12):6063.

31. McKerrow JH (2018) Update on drug development targeting parasite cysteine proteases. PLoS neglected tropical diseases 12(8):e0005850–e0005850.

32. Vuong W, et al. (2020) Feline coronavirus drug inhibits the main protease of SARS-CoV-2 and blocks virus replication. Nature Communications 11(1):4282.

33. Schneider CA, Rasband WS, & Eliceiri KW (2012) NIH Image to ImageJ: 25 years of image analysis. Nature Methods 9(7):671–675.

34. Thoms M, et al. (2020) Structural basis for translational shutdown and immune evasion by the Nsp1 protein of SARS-CoV-2. Science 369(6508):1249.

35. Cox J, et al. (2014) Accurate proteome-wide label-free quantification by delayed normalization and maximal peptide ratio extraction, termed MaxLFQ. Molecular & cellular proteomics: MCP 13(9):2513–2526.

36. Jaimes JA, André NM, Chappie JS, Millet JK, & Whittaker GR (2020) Phylogenetic Analysis and Structural Modeling of SARS-CoV-2 Spike Protein Reveals an Evolutionary Distinct and Proteolytically Sensitive Activation Loop. Journal of molecular biology 432(10):3309–3325.

37. Walls AC, et al. (2020) Structure, Function, and Antigenicity of the SARS-CoV-2 Spike Glycoprotein. Cell 181(2):281–292.e286.

38. Tang T, et al. (2020) Proteolytic activation of the SARS-CoV-2 spike S1/S2 site: a re-evaluation of furin cleavage. bioRxiv:2020.2010.2004.325522.

39. Park J-E, et al. (2016) Proteolytic processing of Middle East respiratory syndrome coronavirus spikes expands virus tropism. Proceedings of the National Academy of Sciences 113(43):12262.

