## Supplementary Appendix for "A cysteine protease inhibitor blocks SARS-CoV-2 infection of human and monkey cells"

### Contents

**Figures** .......... **2-11**

**Methods** .......... **12-27**

**References** .......... **28**

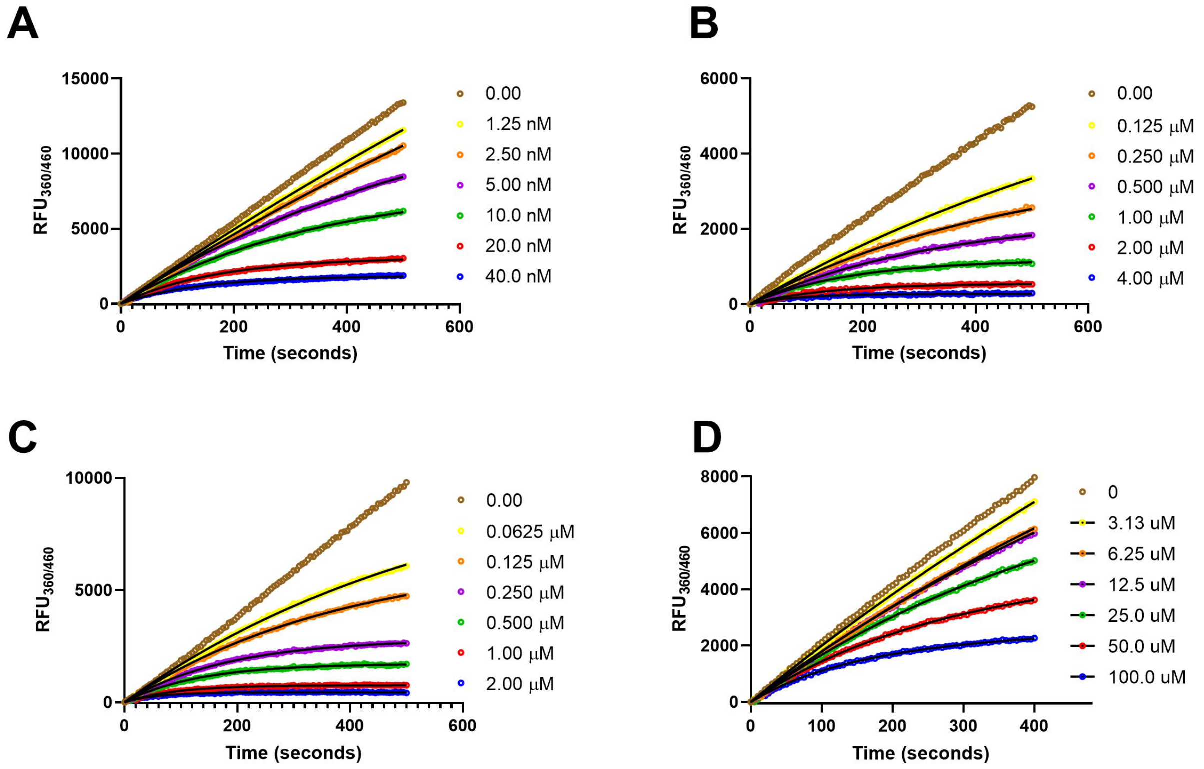


**Fig. S1. Inactivation of cathepsins with K777**. Cathepsin activity monitored by hydrolysis of dipeptide-AMC substrate. Shown are representative plots of inactivation: A) cathepsin L with Z-Phe-Arg-AMC substrate, B) cathepsin B with Z-Phe-Arg-AMC substrate, C) cathepsin K with Z-Leu-Phe-AMC substrate, D) cathepsin C with $NH_2$-Gly-Phe-AMC substrate. Concentration of K777 shown on the figure legends, black lines represent the model of time-dependent inactivation. (eq 1).

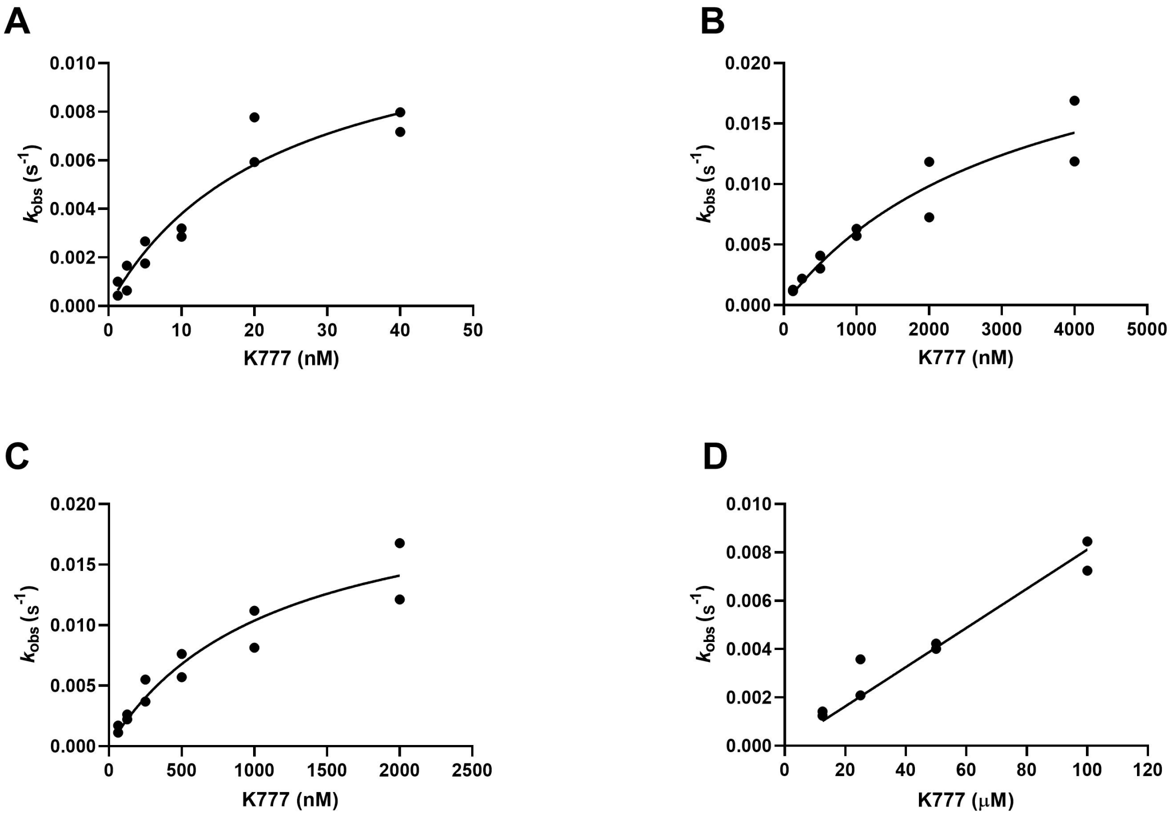


**Fig. S2. Replot of $k_{inact}$ values as a function of K777 concentration**. Data shown are the individual determinations from replicate experiments shown in S2. For **A-C** data was fit to eq 3, for **D** the data was fit to eq 4.

A

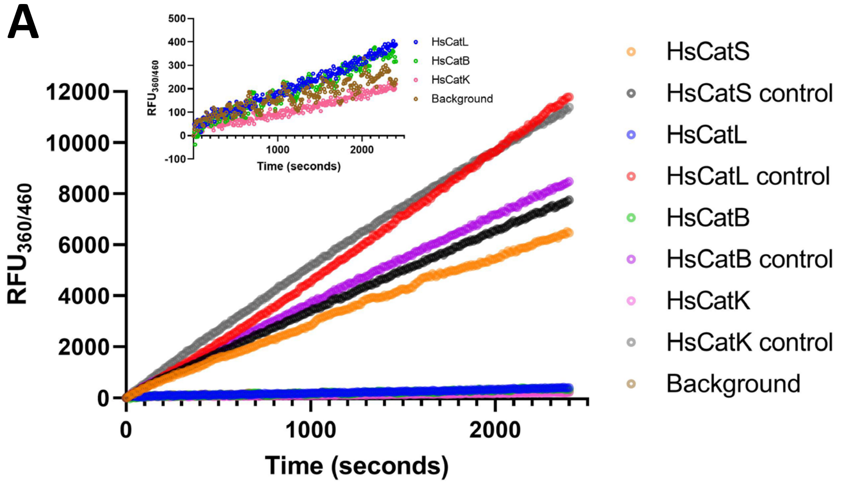


B

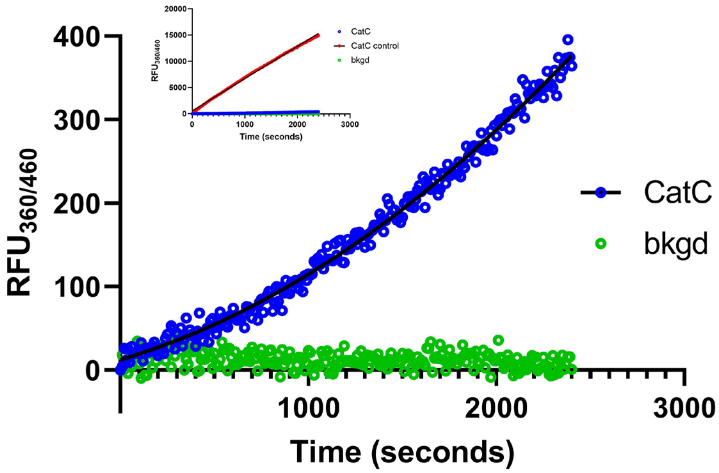


**Fig. S3. Jump-dilution experiment of inactivated cathepsins**. Controls represent enzyme treated with DMSO instead of K777. Background is treated identically to controls except enzyme is replaced with assay buffer. **A** Activity of cathepsins B,K,L, and S following incubation with K777. Inset shows rescaling of the Y-axis and omission of the controls. **B** Activity of cathepsin C following incubation with K777. Inset shows rescaling of the Y-axis and controls. Black line represents model of reactivation, eq 2.

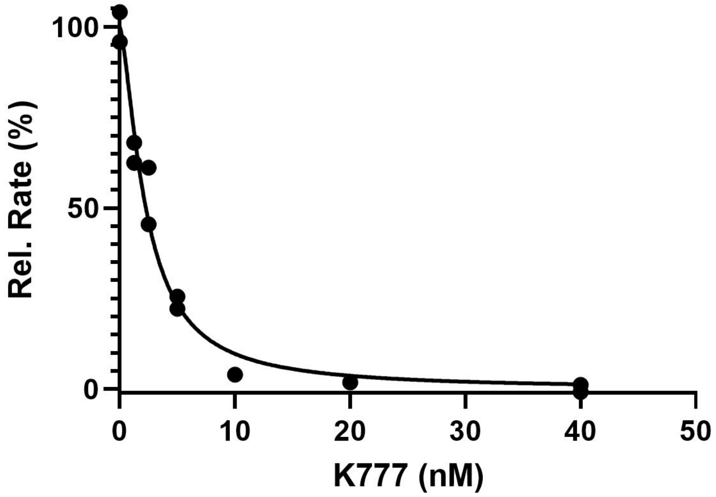


**Fig. S4. Inhibition of cathepsin S with K777**. Inhibition was determined by comparing the steady-state rate at concentrations of K777 (1.25 - 40.0 nM) to DMSO control. Data are plotted from two replicate experiments and was fit to eq. 5

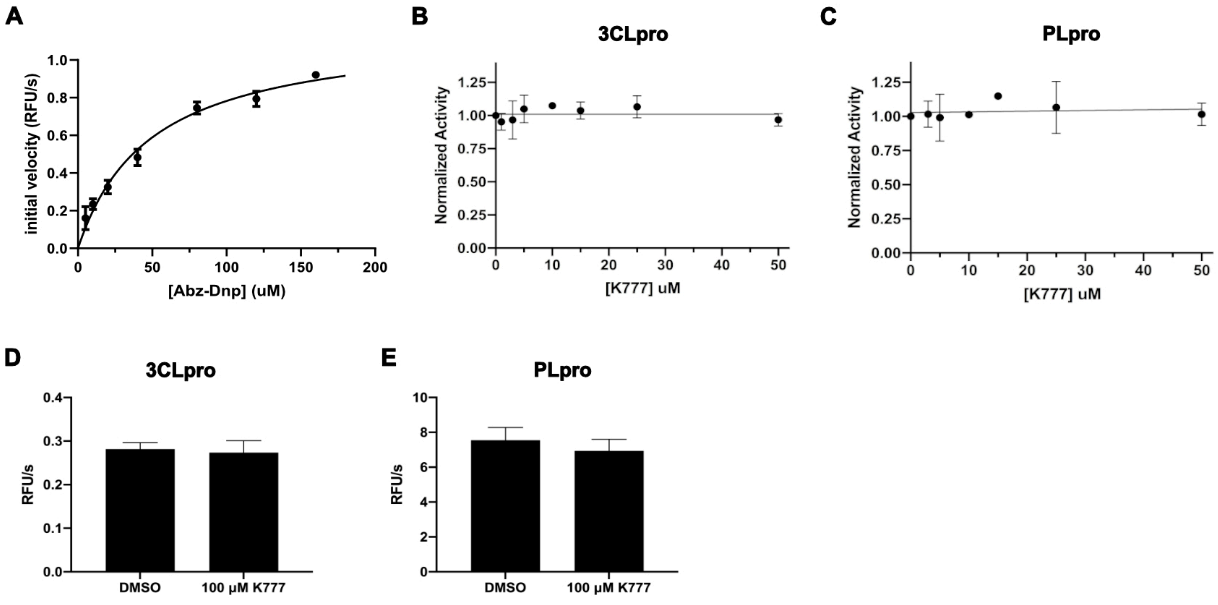


**Fig. S5. A**. Michaelis-Menten kinetic analysis of 3CLpro with the synthetic FRET substrate Abz-Ser-Ala-Val-Leu-Gln*Ser-Gly-Phe-Arg-Lys-(DNP)-$NH_2$. Fitting of this data provided the kinetic parameters: $K_m$= 66 ± 9 µM, $k_{cat}$= 4.9 ± 0.4 $s^{-1}$ and $k_{cat}/K_m$ = 74,000 $M^{-1}s^{-1}$. **B** Average rates determined upon incubation of 25 nM 3CLpro in the presence of 50 µ**M** of Abz-Ser-Ala-Val-Leu-Gln*Ser-Gly-Phe-Arg-Lys-(DNP)-$NH_2$ with varied concentrations of K777 (1-50 µM) at 25°C for 30 minutes. **C.** Average rates determined upon incubation of 10 nM PLpro in the presence of the substrate Arg-Leu-Arg-Gly-Gly-AMC with varied concentrations of K777 (1-50 µM) at 25°C for 30 minutes. **D.** 200 nM of 3CLpro was preincubated with 200 µM of K777 for 30 min and then diluted 2-fold in assay buffer containing 200 µM of Mu-HSSKLQ-AMC. The final assay reaction consisted of 100 nM 3CLpro, 100 µM K777 and 100 µM Mu-HSSKLQ-AMC. **E.** 48.92 nM of PLpro was preincubated with 200 µM of K777 for 30 min and then diluted 2-fold in assay buffer containing 100 µM of Z-RLRGG-AMC. The final assay reaction consisted of 24.46 nM PLpro, 100 µM K777 and 50 µM Mu-HSSKLQ-AMC.

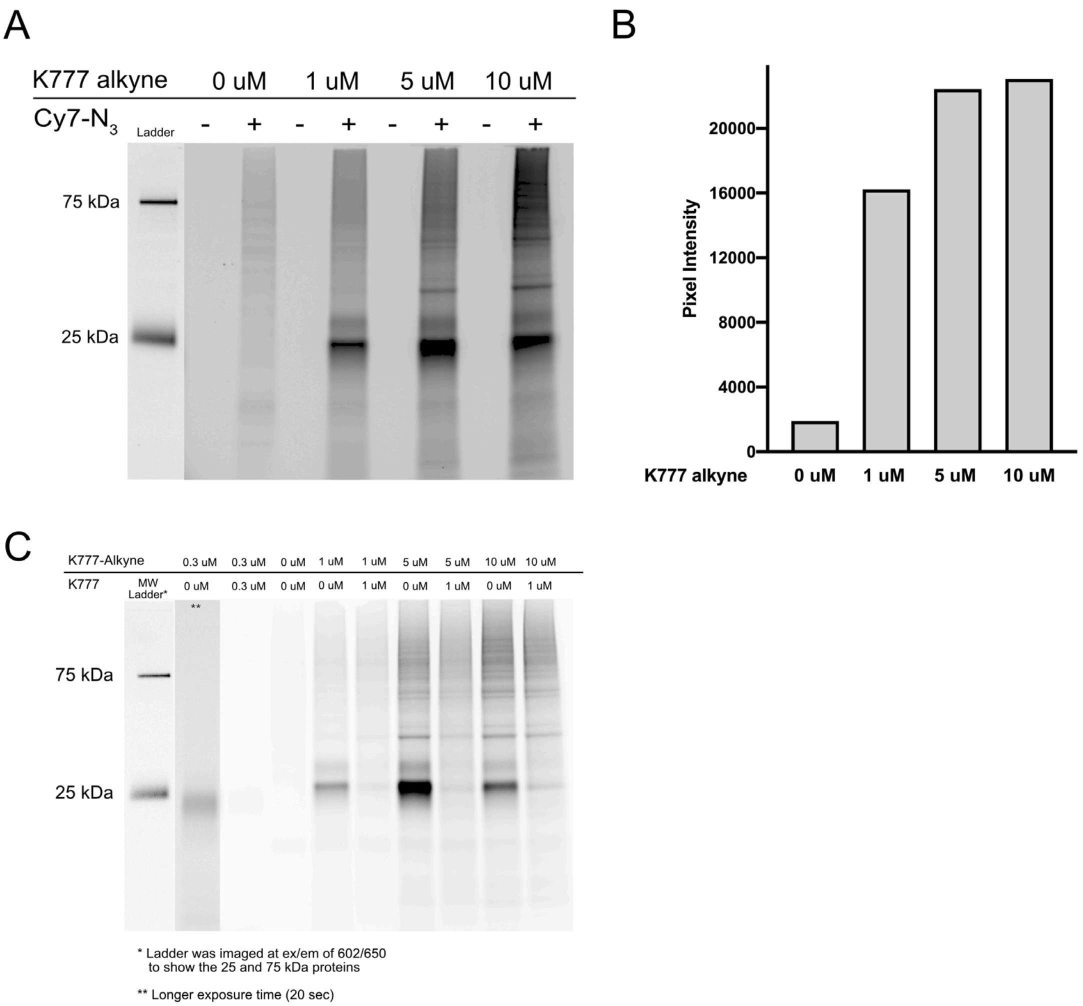


**Fig. S6. A**. In gel fluorescence of non-infected Vero E6 cells treated with varied concentrations of K777 alkyne followed by labeling with Cy7 azide. **B** Densitometry analysis of **A**. **C** In gel fluorescence of non-SARS-CoV-2 infected Vero E6 cells treated with varied concentrations of K777 alkyne and K777.

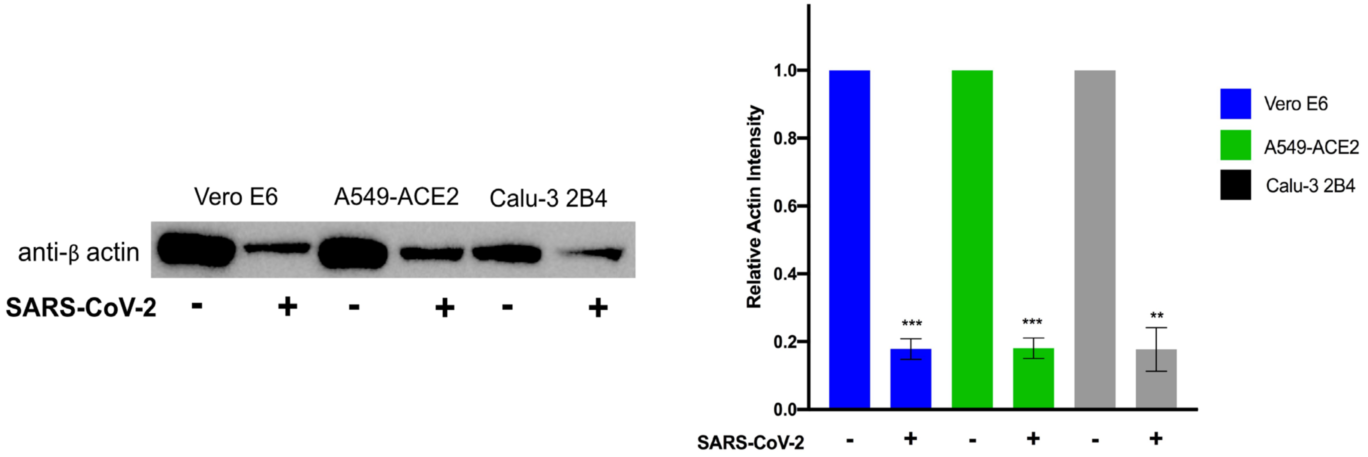


**Fig. S7. Analysis of actin levels in SARS-CoV-2 infected cell lysates.** A. Representative blot of actin levels in virally and non-virally infected cells. B. Densitometry analysis of replicate anti-actin blotting experiments. Signal is normalized to the actin content in cells not infected with SARS-CoV-2.

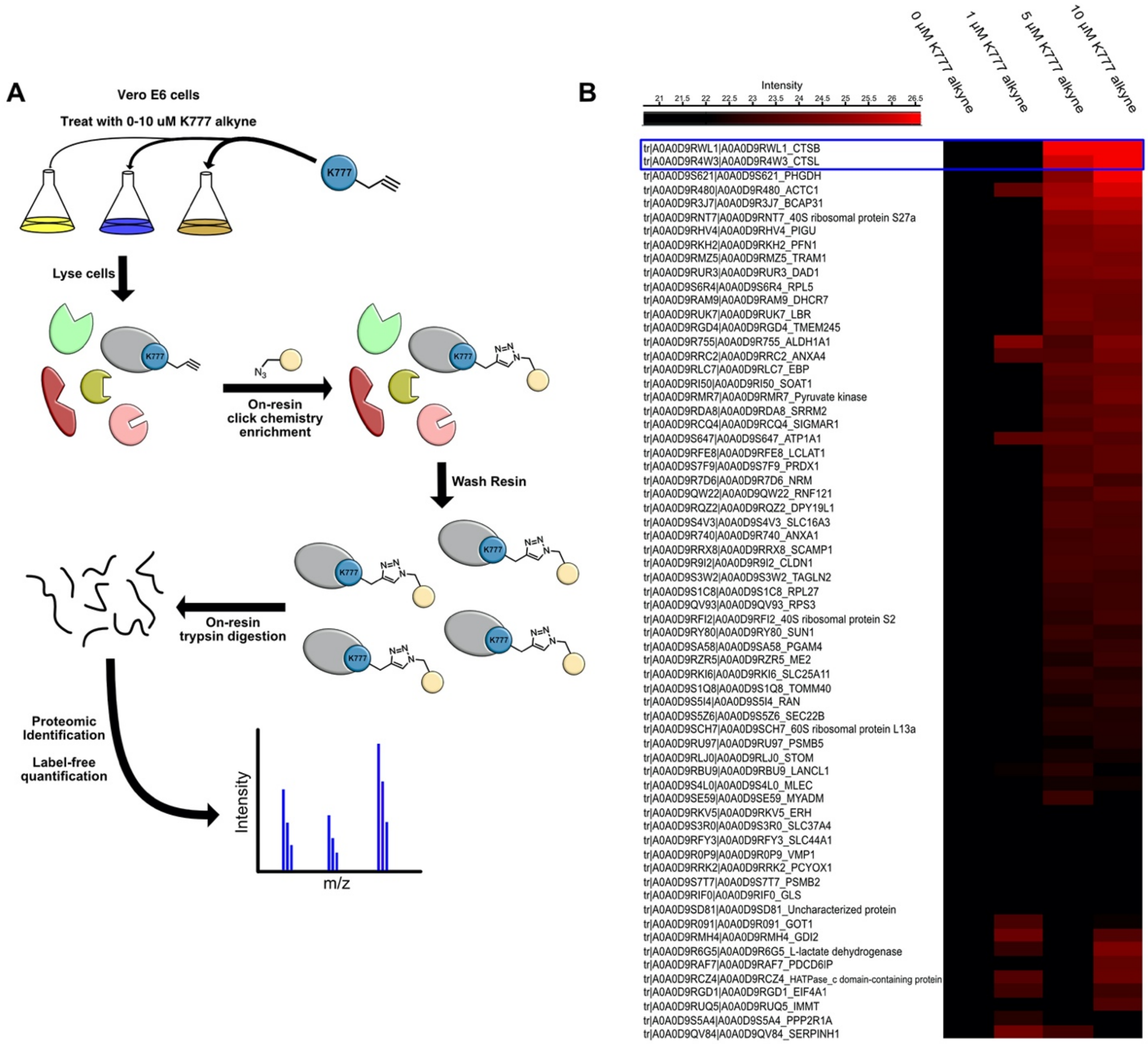


**Fig. S8. Cathepsins B and L are strongly enriched upon K777 alkyne treatment. A.** Schematic of enrichment strategy employed to prepare samples for proteomic processing. **B**. Heat map of proteomic data clustered from highest to lowest intensity. Intensity reports on the summation of the peptide intensities within a particular protein group.

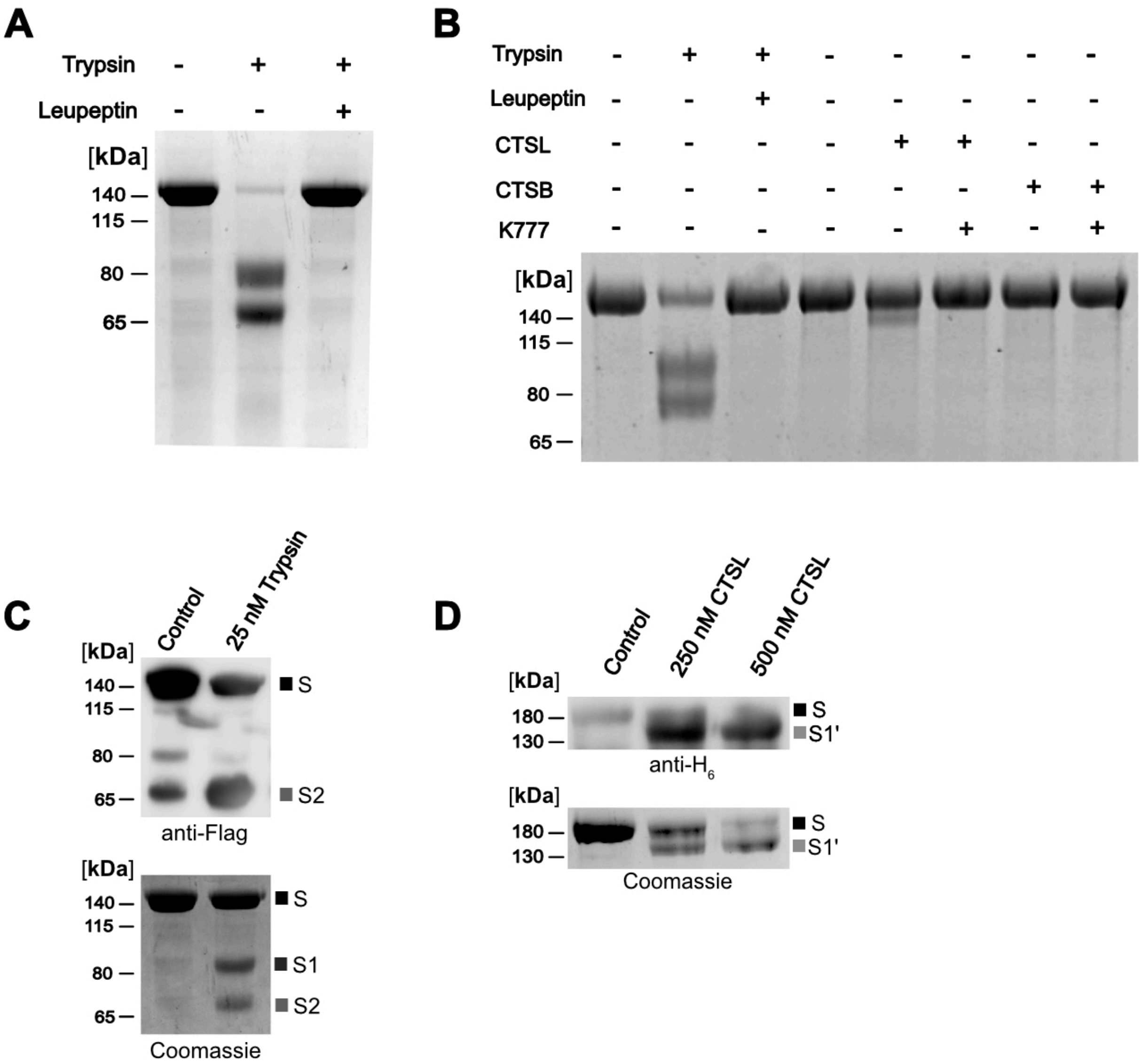


**Fig. S9. Processing of SARS-CoV-2 spike protein by trypsin, CSTB, and CTSL. A.** Trypsin proteolysis of SARS-CoV-2 S produced ~80 kDa and ~65 kDa fragments, which can be blocked by leupeptin. **B** Analysis of processing of the trimeric SARS-CoV-2 S protein by trypsin (25 nM), CTSB (250 nM), and CTSL (250 nM) in the presence and absence of inhibitors. **C.** Representative immunoblotting of trypsin treated SARS-CoV-2 S demonstrates cleavage at the S1/S2 site, which results in signal for the full length S protein and a ~65 kDa fragment corresponding to the cleaved S2 domain. **D.** Immunoblotting of demonstrates that the S1' product produced contains a C-terminal $H_6$ tag, suggestive of CTSL processing in the S1 domain of the protein.

**A**

```
Human CTSL    MNPTLILAAFCLGIASATLTFDHSLEAQWTKWKAMHNRLYGMNEEGWRRAVWEKNMKMIE    60
Monkey CTSL   MNPTFILAALCLGIASATLTFNHSLEAQWTKWKAMHNRLYGMNEEGWRRAVWEKNMKMIE    60
              ****:****:***********:**************************************

Human CTSL    LHNQEYREGKHSFTMAMNAFGDMTSEEFRQVMNGFQNRKPRKGKVFQEPLFYEAPRSVDW    120
Monkey CTSL   LHNQEYSQGKHSFTMAMNTFGDMTSEEFRQVMNGFQNRKPRKGKVFQEPLFYEAPRSVDW    120
              ****** :**********:*****************************************

Human CTSL    REKGYVTPVKNQGQCGSCWAFSATGALEGQMFRKTGRLISLSEQNLVDCSGPQGNEGCNG    180
Monkey CTSL   REKGYVTPVKNQGQCGSCWAFSATGALEGQMFRKTGKLVSLSEQNLVDCSGPQGNEGCNG    180
              ************************************:*:*********************

Human CTSL    GLMDYAFQYVQDNGGLDSEESYPYEATEESCKYNPKYSVANDTGFVDIPKQEKALMKAVA    240
Monkey CTSL   GLMDYAFQYVADNGGLDSEESYPYEATEESCKYNPEYSVANDTGFVDIPKQEKALMKAVA    240
              ********** ************************:************************

Human CTSL    TVGPISVAIDAGHESFLFYKEGIYFEPDCSSEDMDHGVLVVGYGFESTESDNNKYWLVKN    300
Monkey CTSL   TVGPISVAIDAGHESFMFYKEGIYFEPDCSSEDMDHGVLVVGYGFESTESDNSKYWLVKN    300
              ****************:***********************************.*******

Human CTSL    SWGEEWGMGGYVKMAKDRRNHCGIASAASYPTV 333
Monkey CTSL   SWGEEWGMGGYIKMAKDRRNHCGIASAASYPTV 333
              ***********:*********************
```

**B**

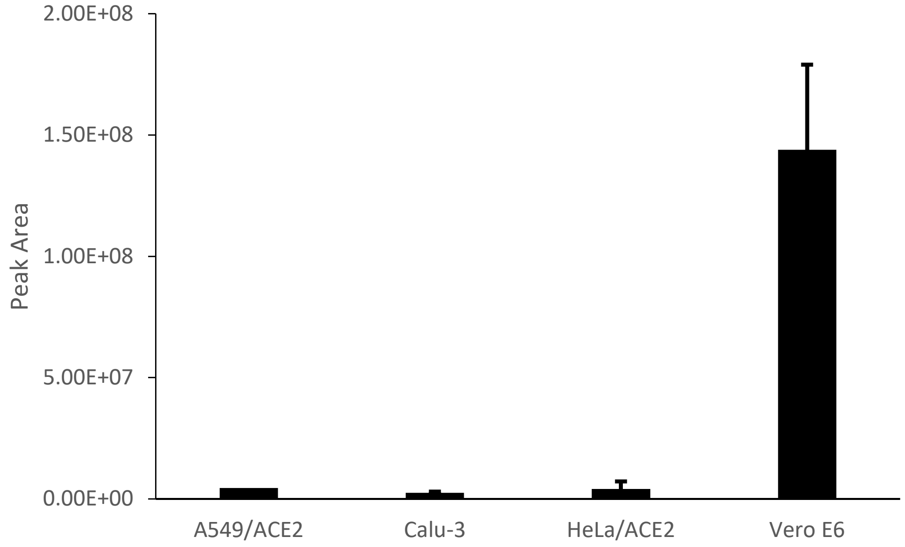


**Fig. S10. Quantitative proteomics of CTSL. A.** Protein alignment of human CTSL and African green monkey CTSL. The C-terminal peptide (NHCGIASAASYPTV) highlight in red font, was found in trypsin-digested protein samples from each cell line and therefore used to compare CTSL between cell lines. **B** Peak area comparison of NHCGIASAASYPTV in four cells lines.

### Supplementary Methods:

#### Expression and purification of 3CLpro: Texas A&M

All genes and plasmids utilized for the expression of 3CLpro were prepared by Genscript and any subsequent subcloning of these genes was verified by sequencing. All chemical reagents for protein purification were obtained from commercial vendors and used without further purification.

*Expression and Purification of 3CL protease-Texas A&M.* Based on the three-dimensional structure of SARS CoV-2 3CLpro (PDB 6Y84.pdb)(1), we designed expression constructs comprising nucleotides 10055-10972 of ORF1AB from GenBank (protein ID QHO62106.1, genome sequence MN988668.1, which encodes 306 amino acids.(2) This gene was *E. coli* optimized and ligated into the into the *Bam*HI and *Xho*I restriction sites of the pGEX-6p1 plasmid (Genscript, USA), resulting in the 3CLpro coding sequence being flanked by an N-terminal GST domain followed by a 3CLpro cleavage sequence (SAVLQ*SGF) and an C-terminal sequence containing a modified PreScission protease sequence (SGVTFQ*GP) preceding a $His_6$ sequence.(3, 4) Upon expression, auto-proteolysis from 3CLpro removed the N-terminal GST tag, yielding the authentic N-terminus (Ser-Gly-Phe). Purification of the processed protein using immobilized nickel affinity chromatography, followed by treatment with HRV 3C protease then generates 3CLpro with authentic N and C termini. This construct was transformed into BL21-DE3 *E. coli* cells for protein expression and a single colony of the transformed cells was used to inoculate a culture of LB broth containing 100 ug/mL ampicillin and was incubated at 37°C overnight. Subsequently, 1 L of LB media containing 100 ug/mL ampicillin was inoculated with the starter culture, and incubated at 37°C until reaching O.D. 600 of 0.6-0.8, at which time expression was induced by the addition of 1 mM isopropyl β-thiogalactoside (IPTG). The cells were allowed to continue growth at 37°C for 4-5 h. and were then harvested by centrifugation (6,300 g at 4°C), and either stored at -80°C or lysed immediately for purification. Cells were suspended in 12 mM Tris-HCl, 120 mM NaCl, 0.1 mM EDTA, 2 mM DTT, pH: 7.5 (Buffer A). The cell slurry was then lysed using either a French press (25,000 psi) or by sonication. Lysates were centrifuged at 26,000g to remove cell debris, and clarified lysates were filtered with a 0.45 µm filter. The filtrates were loaded onto a HisTrap HP column (GE Healthcare), and washed with Buffer A. This was directly followed by elution using a linear gradient to 35% buffer B (12 mM Tris-HCl, 120 mM NaCl, 500 mM imidazole, 0.1 mM EDTA, 2 mM DTT, pH: 7.5) over 25 column volumes. The fractions containing pure 3CLpro-PreSision site-$His_6$ protein, as determined by SDS-PAGE, were pooled, and the protein concentration was determined using the presumed monomeric molecular weight of 33.9 kDa and an extinction coefficient of 32,800 $M^{-1}$ $cm^{-1}$. 3CLpro was twice dialyzed against buffer A at 4°C. Proteolysis of the C-terminal $H_6$ tag of the auto processed translated gene product from the pGEX-6p1 plasmid (3CLpro-HRV 3C protease-$His_6$) was conducted by incubating 3.5 units of Pierce™ HRV 3C Protease

(Thermo Fisher Scientific) per mg of 3CLpro overnight at 4°C overnight in buffer A. Subsequently, the protein mixture was successively loaded onto a 5 mL GSTrap HP column and a 5-mL HisTrap HP column (GE Healthcare), to remove the GST-fused HRV 3C protease and undigested $H_6$ tagged protein. The flow through was collected, analyzed by SDS-PAGE, and pure fractions of the tagless 3CLpro were pooled and concentrated (10 kDa molecular weight cutoff filter, GE Healthcare). The protein was deemed to be ≥95% pure by SDS-PAGE, and was stored at – 80°C in 12 mM Tris-HCl, 120 mM NaCl, 0.1 mM EDTA, 2 mM DTT, (pH 7.5) with 50% glycerol (v/v). Analytical gel filtration using a Superdex 200 Increase 10/300 GL column (Buffer A at a flow rate of 0.7 mL/min), indicated that native 3CLpro was the expected homodimer.

*Recombinant SARS-CoV-2 Mpro protease- UCSD***.** The SARS-CoV-2 Mpro plasmid was provided by Rolf Hilgenfeld (3) and transformed into *E.coli* strain BL21-Gold (DE3). General procedures for the expression, purification, and characterization of **Mpro protease** from SARS CoV-2 are as described(24). Briefly, following *E. coli* culture incubation, cells were lysed by homogenization under native conditions (20 mM Tris-HCl, 150 mM NaCl, 0.25 mM DTT, 5 % glycerol; pH 7.8). Buffer and insoluble protein removed by centrifugation at max speed, 20 min., 4°C . Protein was filtered through 0.45 µM filter . Soluble Mpro was purified by FPLC from the bacterial lysate using $Ni^{2+}$ chelating chromatography (HisTrap FF column, GE Healthcare Life Sciences) under native conditions (20 mM Tris-HCl, 150 mM NaCl, 0.1 mM DTT, 5 % glycerol; pH 7.8). The bound Mpro was eluted using a linear gradient of 0–0.5 M imidazole. The active Mpro fractions were pooled and buffer-exchanged into 20 mM Tris-HCl, 150 mM NaCl, <5 mM imidazole, 1 mM DTT, 5 % glycerol; pH 7.8 using an Amicon Ultracel-10k ultrafiltration device (Millipore) and mixed and incubated with 250 IU of PreScission protease (GenScript) at 4°C overnight, resulting in the target protein with authentic N- and C-termini. The PreScission-treated Mpro was loaded on a tandem of GSTTrap FF (GE Healthcare Life Sciences) and HisTrap FF column (GE Healthcare Life Sciences). Active fractions from the flow through were pooled and stored in 20 mM Tris-HCl, 150 mM NaCl, 1 mM DTT, 5 %. glycerol; pH 8.0. Purification was monitored by a kinetic assay using the peptidyl fluorogenic substrate, morpholinoureidyl (Mu)-His-Ser-Ser-Lys-Leu-Gln-7-amino-4-methylcoumarin (AMC), and by SDS-PAGE. The obtained yield was approximately 6.5 mg of Mpro from 1 L of culture medium.

**Fitting of Kinetic Data**

For cathepsins that were irreversibly inactivated by K777 the data was fit to:

$$Product = \frac{V_o}{k}(1 - e^{-k*t}) \quad \text{(eq 1)}$$

where $V_o$ is the initial rate, $k$ is the observed rate of inactivation, and $t$ is time.

For cathepsins that were reversibly inactivated by K777 the data was fit to:

$$Product = V_s t + \frac{V_o - V_s}{k}(1 - e^{-k*t})\ (eq\ 2)$$

where $V_s$ is the steady-state rate.

The observed rates of inactivation were then replot as a function of K777 concentration and fit to:

$$k_{obs} = k_{react} + \frac{k_{inact}*I}{\left(K_I\left(1+\frac{S}{K_M}\right)+I\right)}\ \text{(eq 3)}$$

where $k_{inact}$ is the intrinsic rate of inactivation, $k_{react}$ is the rate of reactivation, I is the concentration of inhibitor: K777, $K_I$ is the dissociation constant, S is the concentration of substrate, and $K_M$ is the Michaelis constant for the substrate. $K_M$ was determined for enzyme-substrate pairs by plotting observed catalytic rate versus substrate concentration and fitting the data to the Michaelis-Menten equation (data not published). For irreversible inhibitors, the rate of reactivation is essentially zero (i.e. $k_{react} = 0$).

When $k_{obs}$ was unable to be saturated, the second-order rate constant was approximated from fitting the data to:

$$k_{obs} = \left(\frac{k_{inact}}{K_I}\right) I + k_{react}\ (eq. 4)$$

$k_{react}$ was determined through fitting the results of the jump-dilution experiment to eq.2.

For when the inactivation was readily reversible, the steady-state rate was plotted against concentration of K777 and fit to:

$$Rel. Activity = \frac{100}{1+\left(\frac{IC_{50}}{I}\right)^H}\ \text{(eq. 5)}$$

where $H$ is the Hill slope.

**In gel fluorescence of K777-alkyne treated VERO E6 cells Click Chemistry**

Labeling of the K777-alkyne treated cells occurred in a stepwise fashion whereby addition of TBTA in 1:4 DMSO: n-butanol (v/v) (100 µM final) was added to $CuSO_4$ in water (1 mM final), followed by the addition of Cy7-azide (Click Chemistry Tools) in DMSO (20 µM final), 24 µg cell lysate, and finally sodium ascorbate in water (2.5 mM final). This reaction was incubated at room temperature for 1.5 hours and quenched by the addition ice-cold methanol (200 µL), chloroform (75 µL), and water (150 µL) followed by vortexing and centrifugation at 15,000xg at 4 ºC for 15 minutes. The top layer of liquid was carefully removed and discarded and the protein precipitate formed between two liquid phases was not disturbed. An additional 1 mL of ice-cold methanol was added into the tube, vortexed, and centrifuged at 15,000xg at 4 ºC for 20 minutes. The supernatant was discarded and this was repeated an additional time. Following removal of the supernatant the protein pellet was allowed to air-dry at room temperature. To the dried pellet 4% SDS buffer (4% SDS (w/v), 50 mM TEA, 150 mM NaCl, pH 7.4) was added and the pellet was sonicated to achieve a clear solution. Then, the solution was added to 4x loading dye (240 mM Tris-HCl, 8% SDS (w/v), 0.004 % bromophenol blue (to avoid fluorescence from the dye), DTT (5 mM final), and MilliQ water was added to a total volume of 24 µL. Samples were heated to 95 ºC for 5 minutes followed by a brief centrifugation at 5,000xg for 1 min. The ladder (Bio-rad, Precision Plus Protein™ Dual Color Standards) was diluted 10,000 fold in 4% SDS buffer and 4x loading dye to avoid strong fluorescence from the ladder during imaging. Samples were analyzed on a 12% Bis-Tris Nu-PAGE gel (Invitrogen). To each well, 20 µg of labeled lysate was loaded and the gel was run at 60 V for 30 minutes followed by 120 V for ~90 minutes. The gels were destained (10% acetic acid, 40% methanol, 50% water) for 1 hour to overnight in the dark followed by imaging the gel on a ChemiDoc imager (Bio-Rad) using the Cy7 blot setting. To visualize the two fluorescent proteins of the ladder we used the rhodamine setting. Densitometry of in gel fluorescence was conducted using ImageJ. The gels were also analyzed by Coommassie Brilliant Blue R-250 staining (Bio-Rad).

**Mass Spectrometry for K777-alkyne enrichment analysis**

nanoLC-MS/MS analysis was performed using an UltiMate 3000 HPLC system (Thermo Scientific) coupled to a Thermo Scientific Orbitrap Fusion™ Tribrid™ mass spectrometer. 1 µl of each sample was injected onto a PepMap100 C18 5 µm trap cartridge (0.3 × 5 mm) followed by an Acclaim PepMap C18 column (0.075 mm × 150 mm, particle size 3 µm, pore size 100 Å) at a flow rate of 30 µl/min. Peptides were eluted at a flow rate of 0.200 µl/min, using 98% water/2% acetonitrile with 0.1% formic acid (A) and 2% water/98% acetonitrile with 0.1% formic acid (B) as mobile phase. The gradient used was as follows: equilibration at 2% B for 5 minutes, ramping up to 45% at 37 minutes, 90% B at 40 to 46 minutes, ramping down to 2% B at 47 minutes, followed by re-equilibration at 2% B until the end of the run (60 minutes).

Following LC separation, samples were introduced into the mass spectrometer by nanoelectrospray ionization at a spray voltage of 2450 V, with the ion transfer tube temperature set to 275 °C. Data were acquired in top speed mode, with the cycle duration set to 3 seconds. Full scan data were acquired in the orbitrap in the 400-1600 m/z range at a resolution of 120,000 at m/z 200. MS/MS data were acquired in the ion trap in rapid scan mode, using an isolation window of 1.6 m/z with HCD at a fixed normalized collision energy of 28%. Dynamic exclusion was set to 60 seconds.

Data were processed using MaxQuant 1.6.14. Label-free quantification was enabled at the default parameters. The green monkey (*Chlorocebus sabaeus*) proteome (UniProt ID UP000029965; 19,229 protein sequences) was used as the target database. MaxQuant result files were analyzed using Perseus 1.6.14. Reverse hits, potential contaminant hits, hits only identified by site and hits for which only one peptide was detected were removed. Only proteins which were not detected in the control were considered. Intensity values were logarithmized. Finally, proteins which were detected in at least two test samples were hierarchically clustered at default parameters (column clustering was disabled) and plotted as a heat map.

**Proteomic analysis of cell line protein extracts**

Protein extracts from Vero-E6, HeLa/ACE2, Calu-3 and A549/ACE2 cell lines were prepared by sonication in 50 mM MES, pH 6.0, containing 1 mM EDTA, 1 µM pepstatin, 100 µM AEBSF and 1 mM DTT in an ice bath. The extracts were cleared by centrifugation (16,000 g at 4°C for 10 min) and the protein concentration was determined by BCA assay (Pierce). 100 µg of each protein lysate was denatured and reduced with solid urea (8 M final) and DTT (5 mM), and incubated at 56°C for 30 min. Iodoacetamide was added to each sample (15 mM final) using a freshly-made stock solution, and samples were incubated in the dark at 22°C for 30 min. Reactions were quenched with an additional 5 mM DTT, then diluted with 50 mM Tris-HCl pH 7.5 to 1 M urea final concentration prior to addition of sequencing-grade trypsin (sequencing grade, Promega V5113, Madison, WI) at 1:50 trypsin/total protein for digestion overnight at 37 °C. Reactions were quenched by adding 10% TFA to acidify the samples to pH <2 and desalted using C18 spin columns. Two micrograms of the extracted peptides were analyzed on a Q Exactive Mass Spectrometer (Thermo) equipped with an Ultimate 3000 HPLC (Thermo). Peptides were separated by reverse phase chromatography on a C18 column (1.7 m bead size, 75 m 20 cm, heated to 65 °C) at a flow rate of 300 nL/min using a 56-min linear gradient from 4% B to 17% B followed by a 20-min gradient from 17% B to 25% B, with solvent A: 0.1% formic acid (Thermo) in water and solvent B: 0.1% formic acid in acetonitrile (Thermo). Survey scans were recorded over a 310 –1250 m/z range at 70,000 resolution at 200 m/z. MS/MS was performed in data-dependent acquisition mode with HCD fragmentation (28 normalized collision energy) on the 20 most intense precursor ions at 17,500 resolution at 200 m/z. Data was processed using PEAKS 8.5 (Bioinformatics Solutions Inc.). MS2 data were searched against their proteome (Uniprot). Fixed modifications of carbamidomethylation of cysteines (57.02146 Da), variable modification of acetylation of protein N termini (42.0106) and oxidation of methionine (15.99492 Da) were specified. A precursor tolerance of 20 ppm and 0.01 Da for MS2 fragments was defined. Data were filtered to 1% peptide and protein level false discovery rates with the target-decoy strategy.

**Synthetic Chemistry: Synthesis and Characterization of 3CLpro peptide substrate and K777:**

*General synthetic considerations.*

The reagents and starting materials used were obtained from commercial vendors and used as received without any purification. Reactions were carried in an inert atmosphere of nitrogen unless otherwise specified. Progress of the reactions were monitored using Thin Layer Chromatography (TLC) and LC-MS analysis, by employing an HPLC-MS (UltiMate 3000 equipped with a diode array coupled to a MSQ Plus Single Quadrupole Mass Spectrometer, ThermoFisher Scientific) using electrospray positive and negative ionization detectors. HPLC conditions used: column: Phenomenex Luna 5 µm C18(2) 100 Å, 4.6 mm, 50 mm, Mobile phase A: water with 0.1% formic acid (v/v). Mobile phase B: MeCN with 0.1% formic acid (v/v). Temperature: 25 °C. Gradient: 0−100% B over 6 min, then a 2 min hold at 100% B. Flow: 1 mL/min. Detection: MS and UV at 254, 280, 214, and 350 nm. $^{1}H/^{13}C$ NMR spectra were obtained in $CDCl_3$, $CD_3OD$, or DMSO-$d_6$ at 400MHz/100MHz at 298 K on a Bruker Avance III NanoBay console with an Ascend magnet. The following abbreviations were utilized to describe peak patterns when appropriate: br = broad, s = singlet, d = doublet, q = quartet, t = triplet, and m = multiplet. The final compounds used for testing in assays and biological studies had purities that were determined to be >95% as evaluated by their proton NMR spectra and/or their HPLC/MS traces based on ultraviolet detection at 254 nm (K777 and K777 alkyne) or 350 nm (Abz FRET peptide).

All reagents and starting materials were obtained from commercial suppliers and used without further purification unless otherwise stated. Solution phase reactions were conducted under an atmosphere of nitrogen at ambient temperature unless otherwise noted. Reaction progress was monitored using thin-layer chromatography and by HPLC–MS (UltiMate 3000 equipped with a diode array coupled to an ISQ EM single quadrupole mass spectrometer, Thermo Fisher Scientific) using electrospray positive and negative ionization detectors. Reported liquid chromatography retention times ($t_R$) were established using the following conditions: column: Phenomenex Luna 5 µm C18(2) 100 Å, 4.6 mm, 50 mm; mobile phase A: water with 0.1% formic acid (v/v); mobile phase B: MeCN with 0.1% formic acid (v/v); temperature: 25 °C; gradient: 0–100% B over 6 min, then a 2 min hold at 100% B; flow: 1 mL $min^{-1}$; and detection: MS and UV at 254, 280, 214, and 350 nm.

Semi-preparative HPLC purification of compounds was performed on a Thermo Fisher Scientific UltiMate 3000 with a single wavelength detector coupled to a fraction collector. Purifications were conducted using the following conditions: column: Phenomenex Luna 5 µm C18(2) 100 Å, 21.2 mm, 250 mm; mobile phase A: water with 0.1% formic acid (v/v); mobile phase B: acetonitrile with 0.1% formic acid (v/v); temperature: room temperature.

*Fmoc-Lys(DNP)-OH*

A round bottom flask charged with Fmoc-Lys-OH (2 g, 5.43 mmol) was purged with $N_2$ gas to yield positive pressure, followed by the addition of 50 mL anhydrous dichloromethane. Subsequently, DIPEA (2.84 mL, 16.3 mmol) was added, followed by the dropwise addition of 1-fluoro-2,4-dinitrobenzene (700 µL, 5.54 mmol). The mixture was allowed to react for 3.5 hours at which time the reaction mixture was washed with 1N HCl (1 time), water (3 times), brine (1 time), dried over $Na_2SO_4$, and concentrated under reduced pressure. The crude product was subjected to flash purification on silica (2% MeOH:DCM), which following concentration under reduced pressure yielded a bright yellow fluffy powder (1.89 g, 65.2% yield). LC-MS $t_R$: 5.99 min, m/z 535.12 [M+1H], $C_{27}H_{26}N_4O_8$ Calcd. 535.18 [M+1H].

*Boc-2-Abz-OH*

A round bottom flask charged with 2-aminobenzoic acid (2 g, 14.58 mmoles) was dissolved in 20 mL of water and the pH was adjusted to 8 by adding 10 N NaOH dropwise. Subsequently, di-tert-butyl dicarbonate (3.5 g, 16.04 mmol) was dissolved in 20 mL of anhydrous THF was added dropwise to the reaction which was allowed to proceed overnight. The organic was removed under reduced pressure the aqueous layer was acidified with 2N HCL and extracted with ethyl acetate 3 times. The organic layer was washed with water (2 times), and brine (2 times) and dried over dried over anhydrous sodium sulfate and removed under reduced pressure yielding an off-white powder. (2.76 g, 80.7% yield) LC-MS $t_R$: 5.36 min, m/z 236.08 [M-1H], $C_{12}H_{15}NO_4$ Calcd. 236.10 [M-1H].

*Fmoc-SAVLQSGFRK(DNP)-NH₂*

To a syringe fitted with a frit, 200 umol of Rink Amide AM resin (Novabiochem, 0.73 mmol/g) was washed with dichloromethane (DCM) and swelled in N,N-dimethylformamide (DMF). The Fmoc resin was deprotected using 20% piperidine in DMF (v/v) for 15 minutes (3 times), followed by five washes with DMF. Coupling of Fmoc-Lys(DNP)-OH was conducted using 3-fold excess of the amino acid versus resin loading (0.6 mmol), (1-Cyano-2-ethoxy-2-oxoethylidenaminooxy)dimethylamino-morpholino-carbenium hexafluorophosphate (COMU) (0.6 mmol), and DIPEA (1.2 mmol) under agitation for 3 hours (2 times). Washing of the resin with DMF, followed by ninhydrin analysis verified the coupling was complete. The resin was then capped by 45 minutes of agitation with 25% acetic anhydride in DMF (v/v) and DIEA (0.3 mmol), followed by DMF washes (5 times). Subsequent synthesis of the peptide occurred in a stepwise fashion: deprotection of the Fmoc protecting group by 20% piperidine in DMF (v/v) for 15 minutes (2 times), DMF wash (5 times), ninhydrin analysis, addition of Fmoc-AA-OH (0.6 mmol), COMU (0.6 mmol), and DIEA (1.2 mmol) (allowed to pre-activate for 2 minutes prior to addition to resin) and couple for 1 hour, wash resin with DMF (5 times), ninhydrin analysis, cap unreacted amine groups with 25% acetic anhydride in DMF (v/v) and DIPEA (0.3 mmol) for 15 minutes, wash resin with DMF (5 times). All steps were carried out under constant agitation. Test cleavage of the peptide was conducted using ~5-10 mg of resin suspended in hexafluoroisopropanol (HFIP) containing 0.5 N HCl (aq), 5% v/v water, and 2.5% v/v triisopropylsilane for two hours. The solution was then removed under a stream of $N_2$ gas and the resulting material was dissolved in acetonitrile, filtered, and analyzed by LC-MS, showing that the peptide was the major product (m/z 740.4 [M+2H]; 1479.7 [M+1H] or 740.4 [M+2H] calcd.). The peptide-laden resin was then washed with DCM (3 times), dried under vacuum, and stored in a desiccator for further synthesis.

*Abz-SAVLQSGFRK(DNP)-NH$_2$*

To a syringe fitted with a frit, half of dry resin bound Fmoc- (DNP)-Rink Amide peptide was suspended in DMF for 20 minutes. Treatment with 20% piperidine in DMF (v/v) for 15 minutes (3 times), followed by DMF washes (5 times) and a ninhydrin test confirmed removal of the Fmoc group. To the free amine, Boc-2-Abz-OH (0.3 mmol), COMU (0.3 mmol), and DIPEA (0.6 mmol) (allowed to pre-activate for 2 minutes) was added and allowed to couple overnight. The resin was washed with DMF (5 times) and the coupling was repeated for another four hours to enhance the yield. The resin was then washed with DMF (5 times), DCM (3 times), and MeOH (3 times). Subsequent cleavage of the peptide with HFIP containing 0.5 N HCl (aq), 5% v/v water, and 2.5% v/v triisopropylsilane for four hours (2 times) was followed by removal of the solution via rotary evaporator and precipitation of the peptide in ice cold diethyl ether. The crude mixture was solubilized in a small volume of DMF and subjected to semi-preparative HPLC (5%-52% B over 20 mins, then 52-100% B to 26 minutes, and 100% B until 29 minutes at 21.2 ml/min). The fractions containing pure peptide (≥95% pure) were then lyophilized to yield a fluffy yellow powder (42.4 mg, 29.2% yield as a formic acid salt) LC-MS $t_R$: 5.36 min, m/z 688.89 [M+2H], 459.79 [M+3H], $C_{61}H_{89}N_{19}O_{18}$ Calcd 1376.66 [M+1H], 688.83 [M+2H], 459.56 [M+3H].

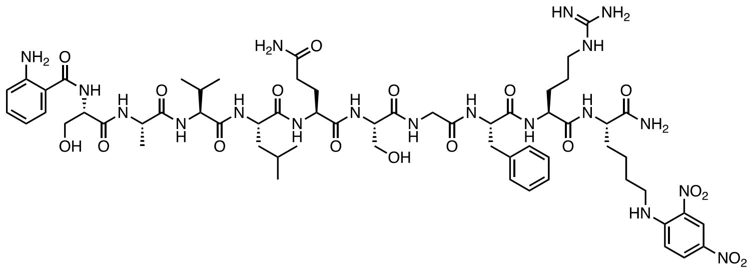

*Abz-SAVLQSGFRK(DNP)-NH₂*

**Synthetic Schemes for K777 and Propargyl-K777[a]:**

**1. Synthesis of K777**

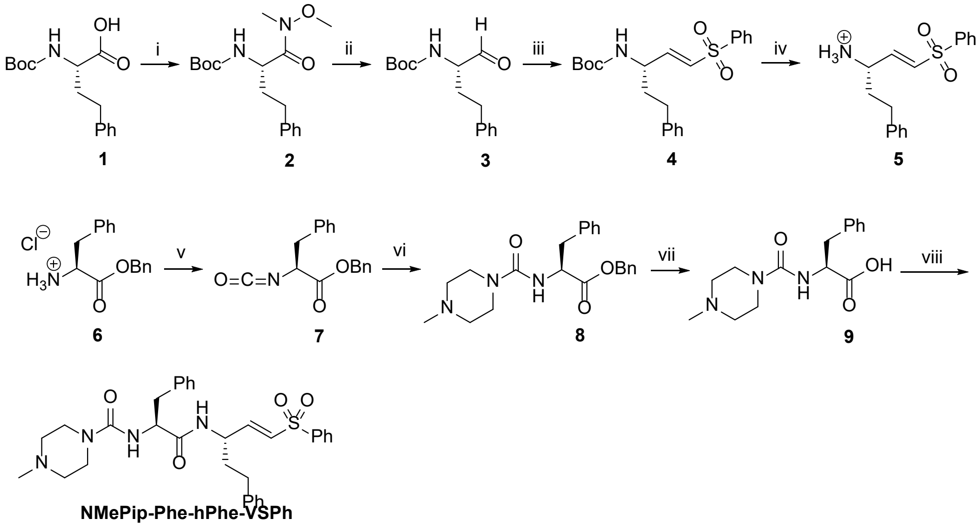

**NMePip-Phe-hPhe-VSPh**

**K777**

**2. Synthesis of Propargyl-K11777**

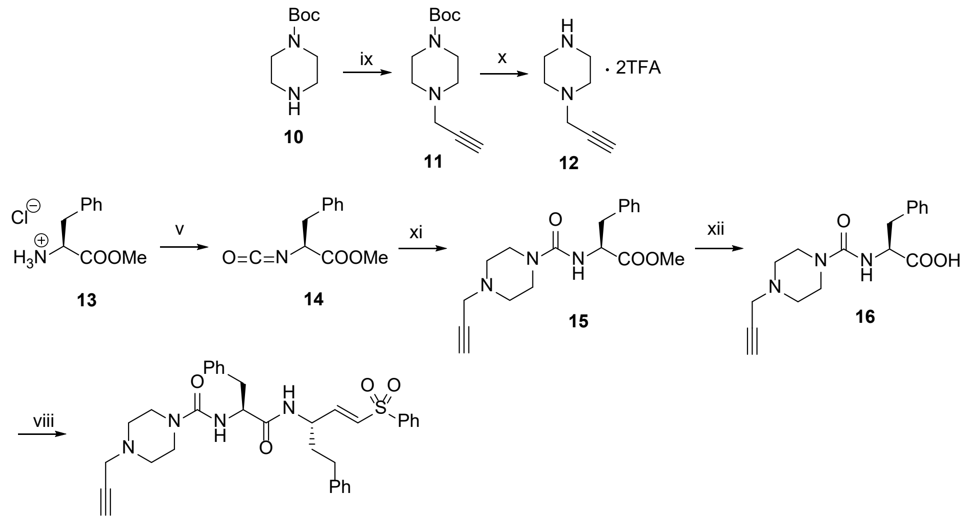

**N(Propargyl)Pip-Phe-hPhe-VSPh**

**Propargyl-K777**

[a]Conditions and reagents: (i) *N,O*-dimethylhydroxylamine hydrochloride, T3P, DIPEA, DCM, 0 ºC, 30 min; (ii) LAH, THF, -10 ºC, 30 min; (iii) Diethyl (phenylsulfonyl)methane phosphonate, LHMDS, THF, -10 ºC, 20 min, 0 ºC, 1 h; (iv) TFA, DCM, 0 ºC, 3h; (v) a) DCM, Sat. aq. $NaHCO_3$, b) triphosgene, 0 ºC, 15 min; (vi) 1-Methylpiperazine, THF, DIPEA, 0 ºC, 10 min, 25ºC, 16 h; (vii) 10% Pd/C, MeOH, 25ºC, 20 h; (viii) **5**, T3P, DIPEA, DCM, -10 ºC, 10 min, 0 ºC, 1 h; (ix) propargyl bromide, DIPEA, $CHCl_3$, -10 ºC, 30 min, 25 ºC, 18 h; (x) TFA, DCM, 0 ºC, 3h, 25 ºC, 1 h; (xi) **12**, THF, DIPEA, 0 ºC, 1 h, 25 ºC, 16 h; (xii) a) LiOH, THF, $H_2O$, 0 ºC, 6 h, b) 4N HCl in dioxane to pH 2.

**Synthesis of K777:**

Boc-protected L-homophenylalanine **1** was purchased from a commercial source, converted to Weinreb amide by T3P-catalyzed coupling to *N*,*O*-dimethylhydroxylamine hydrochloride to afford **2**. Reduction of the Weinreb amide **2** using LAH at −10 °C in anhydrous THF provided the Boc-L-homophenylalanine aldehyde **3**. Horner−Wadsworth−Emmons reaction of the aldehyde **3** with the commercially available Diethyl (phenylsulfonyl)methane phosphonate yielded the vinyl phenyl sulfone **4**.

L-Phenylalanine benzyl ester hydrochloride **6** was reacted with triphosgene to give Isocyanate **7**, which was further reacted with 1-Methylpiperazine to give NMe-Pip-Phe-OBn **8**. NMe-Pip-Phe-OBn was subjected to debenzylation using 10% Pd/C with $H_2$ gas to give NMe-Pip-Phe-OH **9**. TFA Deprotection of the Boc group of the vinyl phenyl sulfone **4** gave the TFA salt **5**. The T3P-catalyzed coupling of **5** with NMe-Pip-Phe-OH **9** gave the required compound **K777**.

**Synthesis of Propargyl-K777:**(5)

Commercially available t-butyl Piperazine-1-carboxylate (1-N-Boc-piperazine) **10** was reacted with Propargyl bromide to give 1-N-Boc-Propargyl-Piperazine **11**, which was further treated with TFA to give 1-N-Propargyl-Piperazine TFA salt **12**.

L-phenylalanine methyl ester hydrochloride was reacted with triphosgene to give Isocyanate **14**, which was further reacted with **12** to give N-Propargyl-Pip-Phe-OMe **15**. The compound **15** was subjected to LiOH hydrolysis to give N-Propargyl-Pip-Phe-OH **16**. T3P-catalyzed coupling of **5** with acid **16** gave the required compound **Propargyl-K777**.

*4-methyl-N-((S)-1-oxo-3-phenyl-1-(((S,E)-5-phenyl-1-(phenylsulfonyl)pent-1-en-3-yl)amino)propan-2-yl)piperazine-1-carboxamide (**K777**, NMePip-Phe-hPhe-VSPh)*. To a suspension of TFA salt **5** (0.100 g, 0.240 mmol) in DCM (5 mL) at -10°C, was added dropwise DIPEA (0.48 mL, 2.744 mmol), followed by addition of the acid **9** (0.100 g, 0.343 mmol) and dropwise addition of T3P (0.33 mL, 0.515 mmol). The reaction was continued at the same temperature for 10 min and 0°C for 1h. Upon the completion of the reaction as revealed by TLC analysis (MeOH/DCM = 1:10, v/v), the reaction mixture was diluted with DCM (50 mL), and then washed with Sat. aq. $NaHCO_3$ (1X), $H_2O$ (3X) and brine (1X). The organic layer was dried over anh. $Na_2SO_4$ and filtered. The filtrate was concentrated *in vacuo* to afford the crude product, which was purified by silica gel column chromatography using a gradient of 1% - 10% of MeOH in DCM as eluent to yield the pure product 4-methyl-N-((*S*)-1-oxo-3-phenyl-1-(((*S*,*E*)-5-phenyl-1-(phenylsulfonyl)pent-1-en-3-yl)amino)propan-2-yl)piperazine-1-carboxamide (***K777***, NMePip-Phe-hPhe-VSPh, White solid, 0.089 g, 0.155 mmol, 45% yield). $^1$H NMR (400 MHz, $CDCl_3$) δ 1.62 – 1.87 (m, 2H), 2.22 (s, 3H), 2.23 – 2.29 (m, 4H), 2.42 – 2.57 (m, 2H), 2.94 – 3.05 (m, 2H), 3.22 – 3.35 (m, 4H), 4.52 – 4.68 (m, 2H), 5.26 (d, 1H, $J$ = 7.8 Hz), 6.12 (dd, 1H, $J_1$ = 1.5 Hz, $J_2$ = 15.1 Hz), 6.78 (dd, 1H, $J_1$ = 5.0 Hz,

$J_2$ = 15.1 Hz), 6.98 (d, 2H, $J$ = 7.0 Hz), 7.06 – 7.30 (m, 9H), 7.51 (t, 2H, $J$ = 7.6 Hz), 7.57 – 7.63 (m, 1H), 7.78 – 7.86 (m, 2H); $^{13}$C NMR (100 MHz, $CDCl_3$) δ 31.8, 35.7, 38.9, 43.8, 46.0, 49.1, 54.5, 56.0,126.2, 127.1, 127.6, 128.4, 128.5, 128.6, 129.3, 129.4, 130.5, 133.5, 136.7, 140.4, 140.5, 145.9, 157.0, 172.1; LC-MS *m/z* 575.35, 576.38 $[M+H]^+$, ($C_{32}H_{38}N_4O_4S^+$ Calcd 575.27); $t_R$ = 3.22 min.

*N-((S)-1-oxo-3-phenyl-1-(((S,E)-5-phenyl-1-(phenylsulfonyl)pent-1-en-3-yl)amino)propan-2-yl)-4-(prop-2-yn-1-yl)piperazine-1-carboxamide (**Propargyl-K**777, N(Propargyl)Pip-Phe-hPhe-VSPh).* Followed the procedure from synthesis of ***K**777*, using TFA salt **5** (1.513 g, 3.640 mmol), DCM (60 mL), acid **16** (1.915 g, 6.072 mmol), DIPEA (8.50 mL, 49 mmol) and T3P (5.80 mL, 9.1 mmol). White solid, 1.553 g, 2.594 mmol, 71% yield. $^1$H NMR (400 MHz, $CDCl_3$) δ 1.59 – 1.87 (m, 2H), 2.21 (t, 1H, $J$ = 2.4 Hz), 2.36 – 2.57 (m, 6H), 2.95 – 3.04 (m, 2H), 3.22 – 3.37 (m, 6H), 4.53 – 4.66 (m, 2H), 5.25 (d, 1H, $J$ = 7.7 Hz), 6.11 (dd, 1H, $J_1$ = 1.5 Hz, $J_2$ = 15.1 Hz), 6.78 (dd, 1H, $J_1$ = 5.0 Hz, $J_2$ = 15.1 Hz), 6.96 – 7.03 (m, 2H), 7.07 – 7.23 (m, 9H), 7.46 – 7.56 (m, 2H), 7.57 – 7.63 (m, 1H), 7.79 – 7.87 (m, 2H); $^{13}$C NMR (100 MHz, $CDCl_3$) δ 31.8, 35.7, 38.7, 43.7, 46.8, 49.1, 51.2, 55.9, 73.6, 78.2, 126.2, 127.1, 127.6, 128.4, 128.5, 128.6, 129.3 (2C), 130.5, 133.4, 136.7, 140.3, 140.5, 145.8, 156.9, 172.0; LC-MS *m/z* 599.25, 600.28 $[M+H]^+$, ($C_{34}H_{38}N_4O_4S^+$ Calcd 599.27); $t_R$ = 3.68 min.

### ABBREVIATIONS USED

DIPEA, *N*,*N*-Diisopropylethylamine; hPhe, homophenylalanine; LAH, lithium aluminum hydride; LHMDS, lithium bis(trimethylsilyl)amide; NMePip, N-methylpiperazinyl; T3P, propylphosphonic anhydride; TFA, trifluoroacetic acid; VSPh, vinyl sulfone phenyl.

**HPLC trace and and MS of *Abz-SAVLQSGFRK(DNP)-NH$_2$***

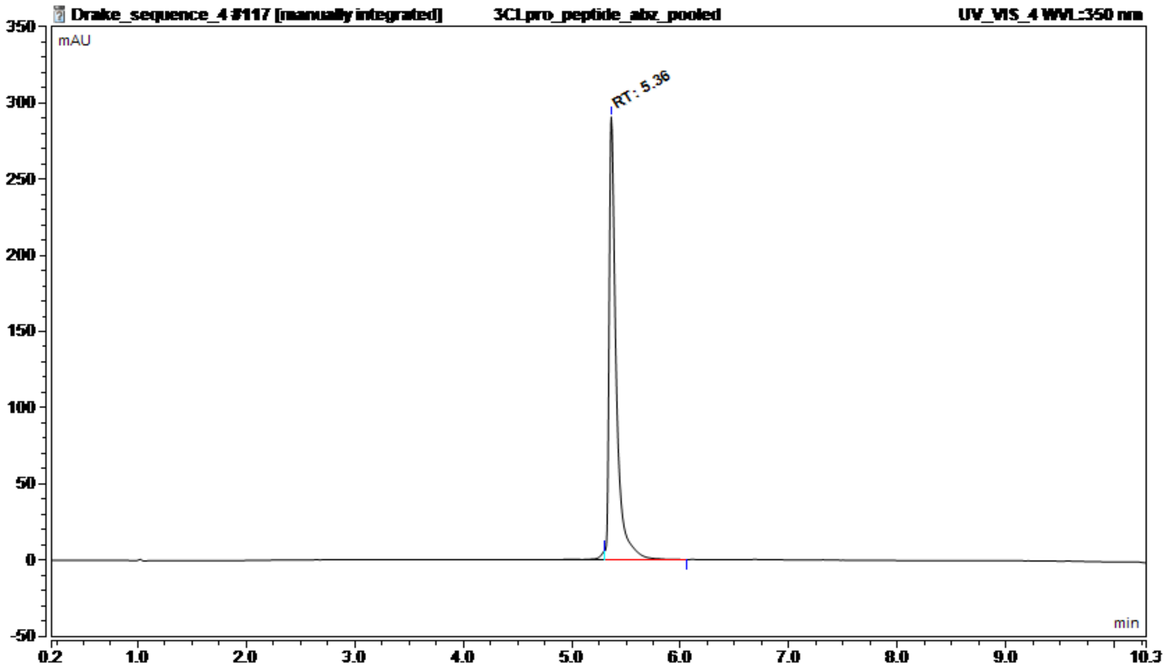


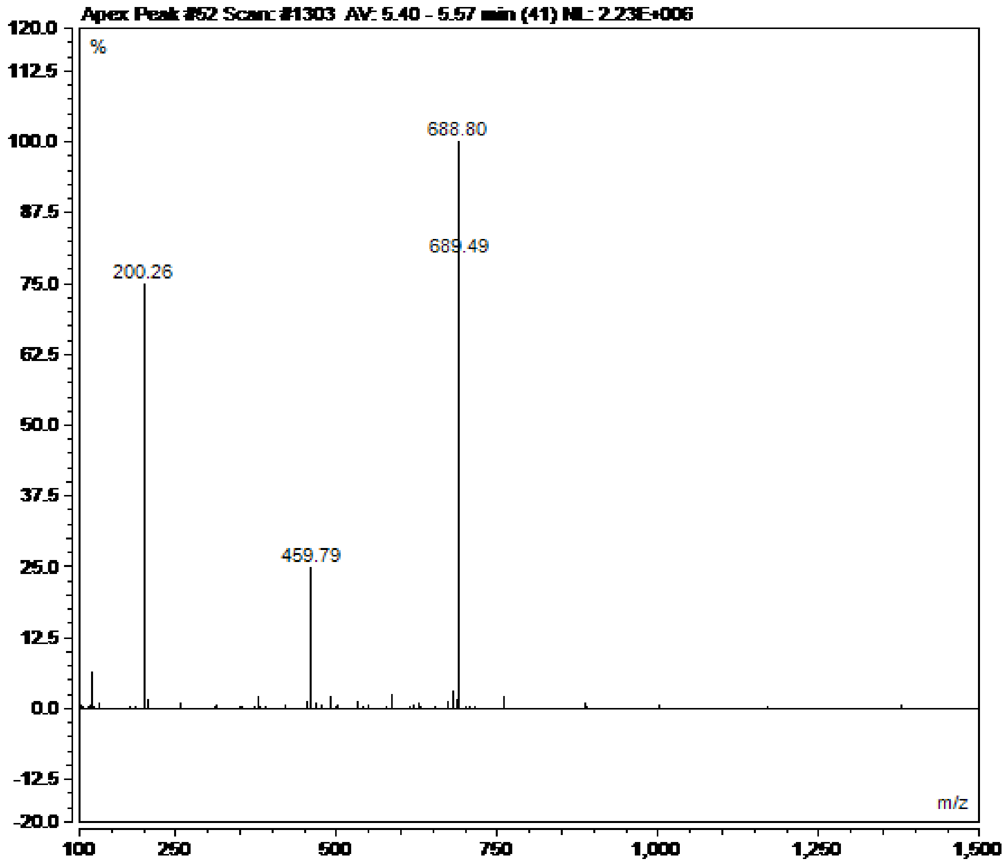

#### HPLC Trace of K777 (NMePip-Phe-hPhe-VSPh)

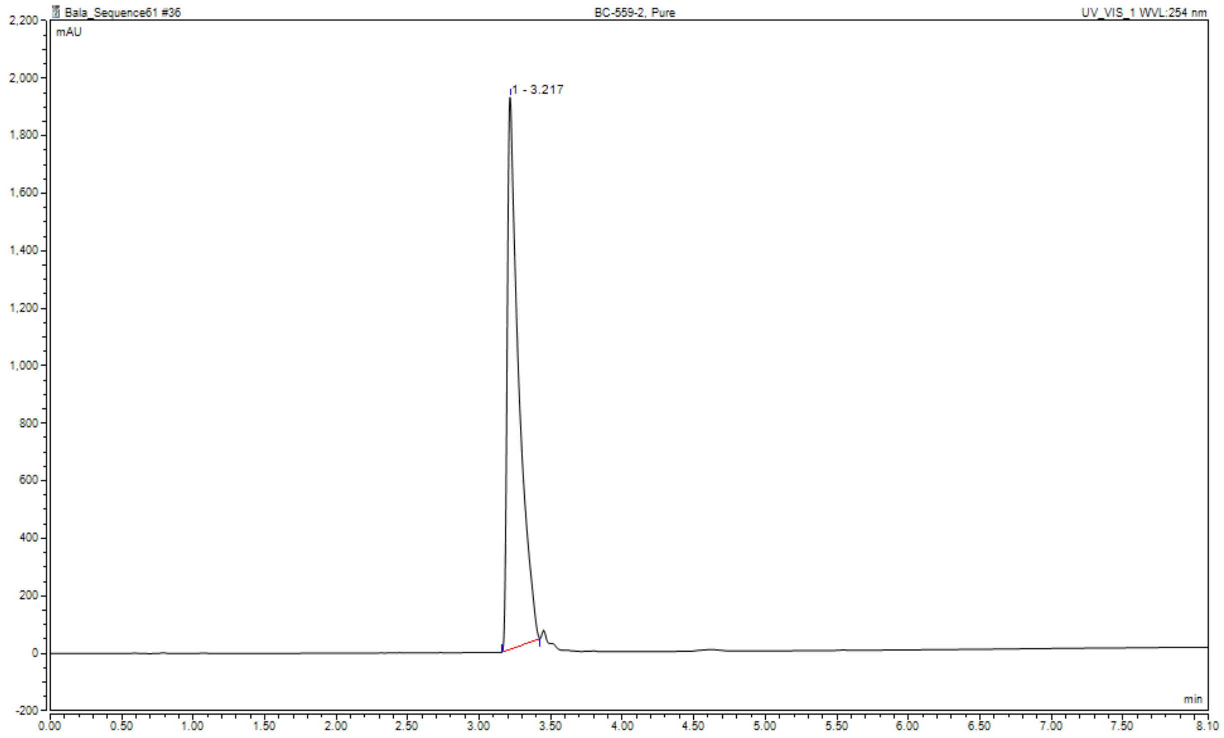


#### HPLC Trace of Propargyl-K777 (N(Propargyl)Pip-Phe-hPhe-VSPh)

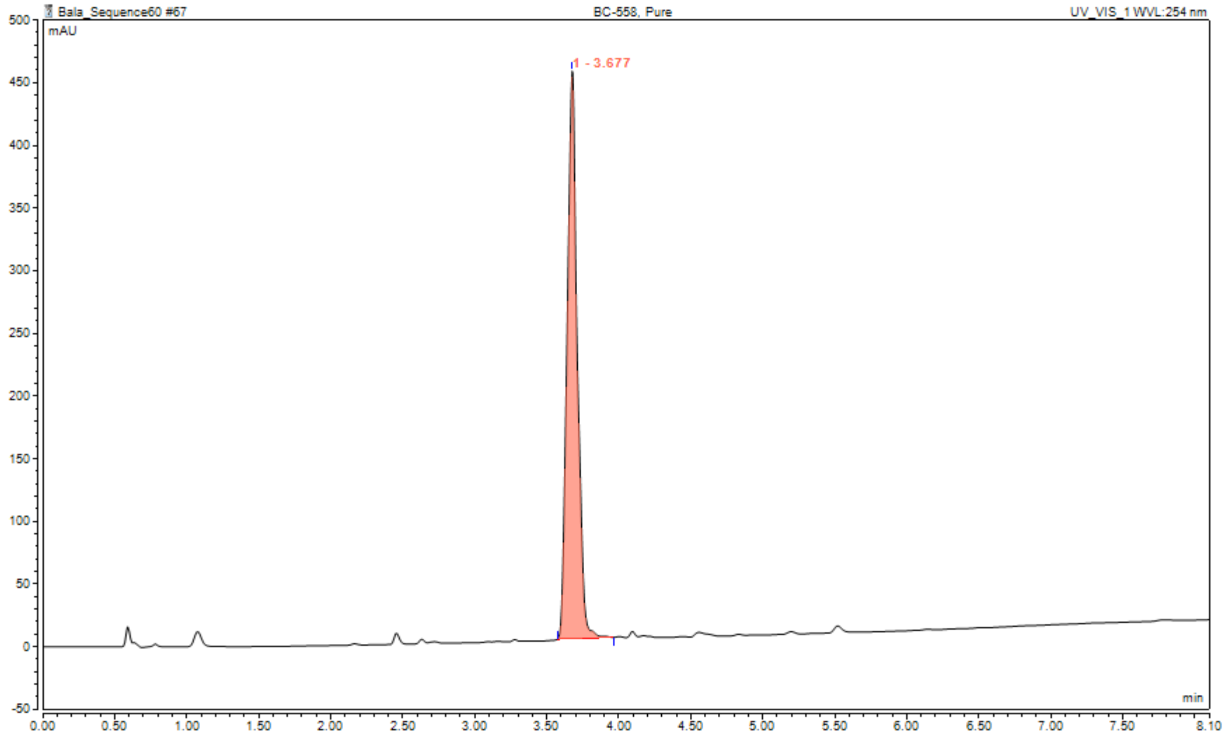

$^{1}$H NMR spectrum of **K777 (NMePip-Phe-hPhe-VSPh)**

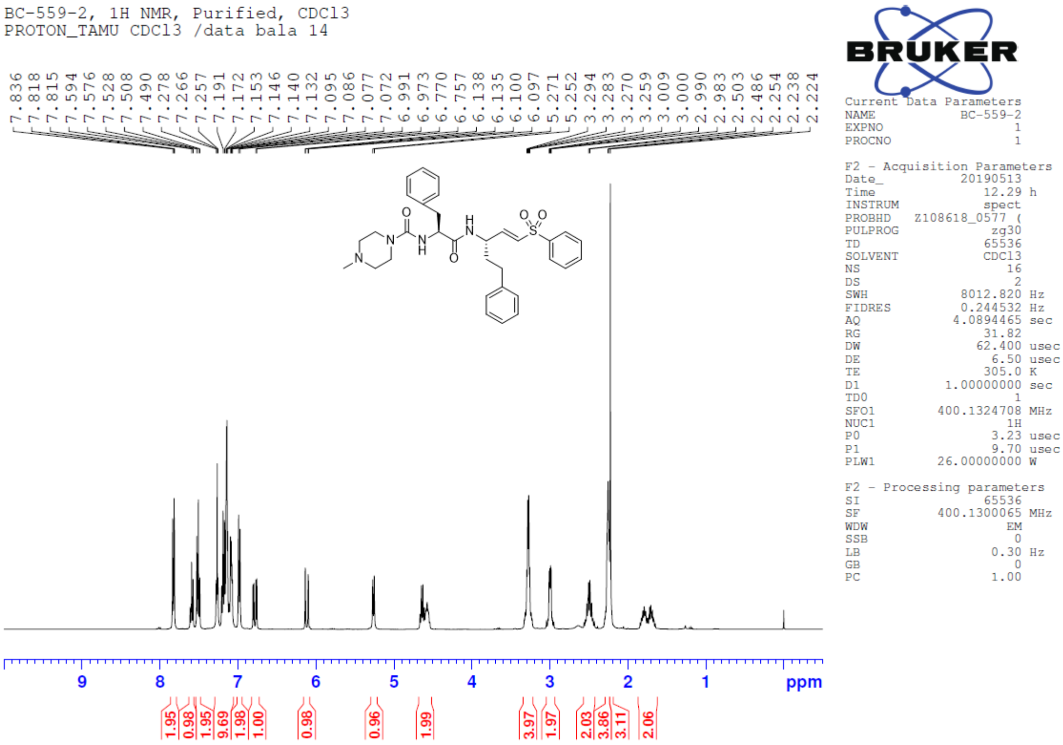


$^{13}$C NMR spectrum of **K777 (NMePip-Phe-hPhe-VSPh)**

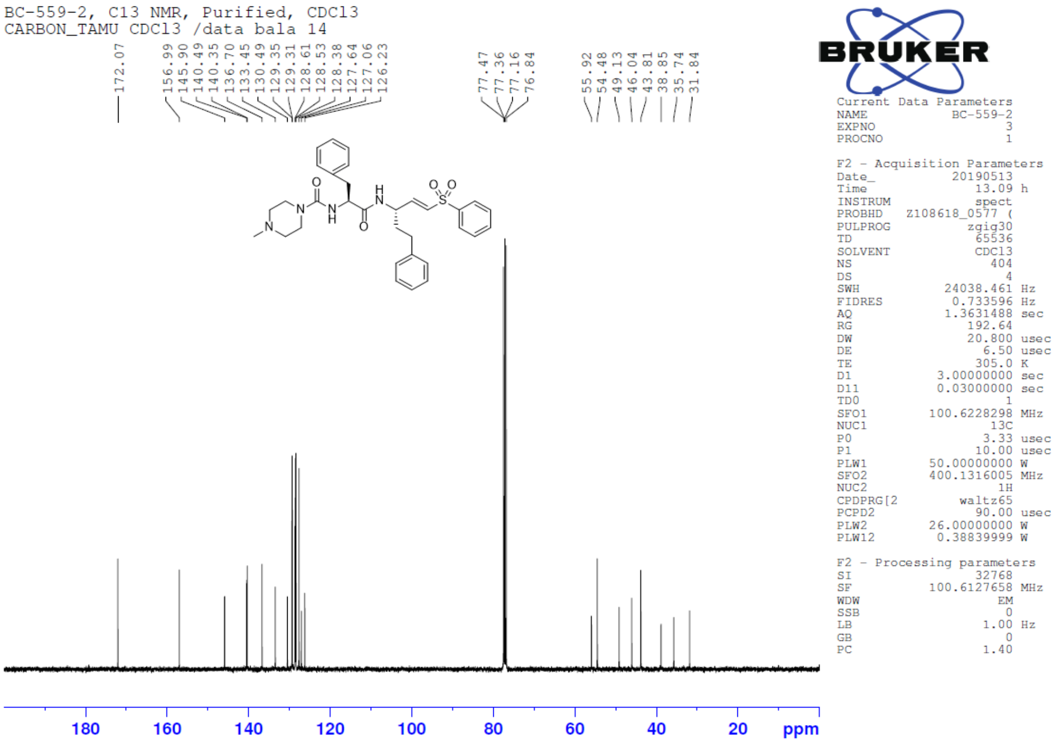

$^{1}H$ NMR spectrum of **Propargyl-K777 (N(Propargyl)Pip-Phe-hPhe-VSPh)**

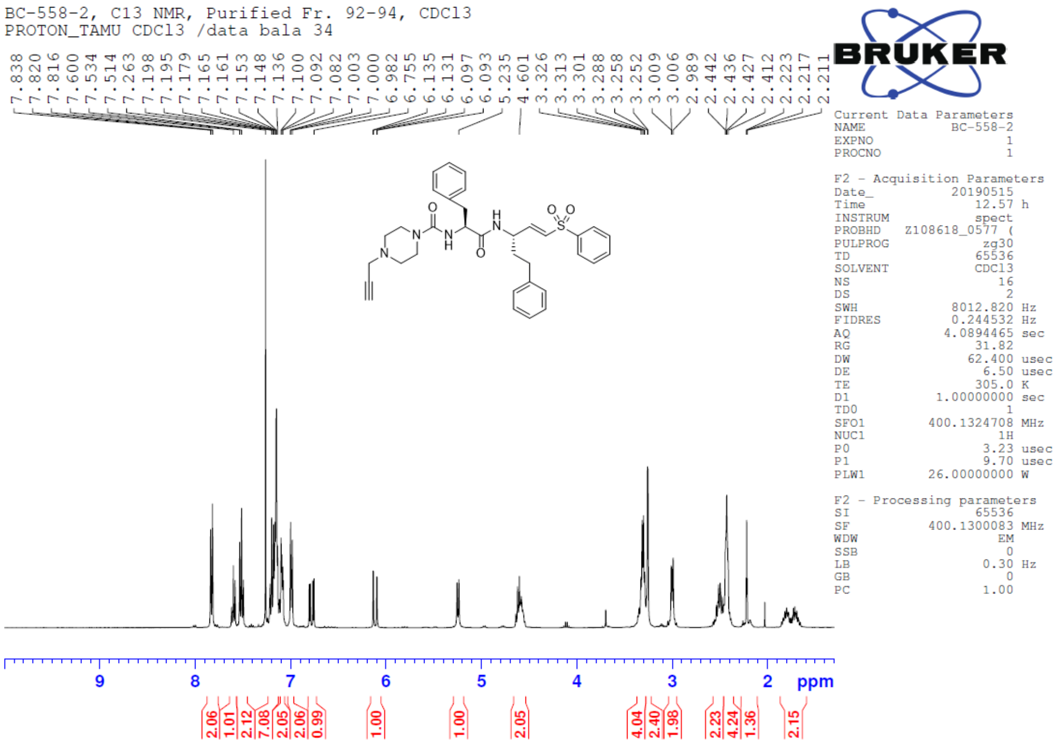


$^{13}C$ NMR spectrum of **Propargyl-K777 (N(Propargyl)Pip-Phe-hPhe-VSPh)**

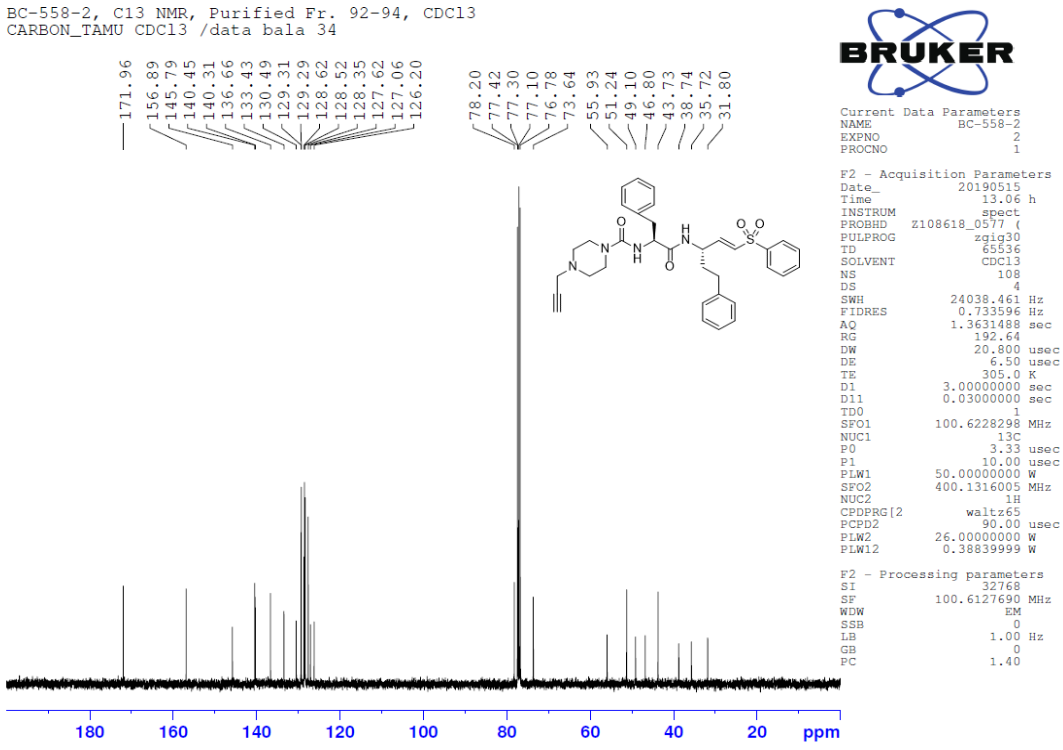

**References:**

1. C. D. Owen CD*, et al.* (2020) 6Y84, SARS-CoV-2 main protease with unliganded active site (2019-nCoV, coronavirus disease 2019, COVID-19). (RCSB Protein Data Bank, RCSB Protein Data Bank).
2. Chen L*, et al.* (2020) RNA based mNGS approach identifies a novel human coronavirus from two individual pneumonia cases in 2019 Wuhan outbreak. *Emerging Microbes & Infections* 9(1):313-319.
3. Zhang L*, et al.* (2020) Crystal structure of SARS-CoV-2 main protease provides a basis for design of improved α-ketoamide inhibitors. *Science* 368(6489):409.
4. Xue X*, et al.* (2007) Production of authentic SARS-CoV M(pro) with enhanced activity: application as a novel tag-cleavage endopeptidase for protein overproduction. *Journal of molecular biology* 366(3):965-975.
5. Yang P-Y, Wang M, He CY, & Yao SQ (2012) Proteomic profiling and potential cellular target identification of K11777, a clinical cysteine protease inhibitor, in Trypanosoma brucei. *Chemical Communications* 48(6):835-837.
