## Supplementary material for "A cysteine protease inhibitor blocks SARS-CoV-2 infection of human and monkey cells": Cover Letter

UC San Diego

October 23, 2020

Dear Editor:

Please find the attached manuscript entitled "**A cysteine protease inhibitor blocks SARS-CoV-2 infection of human and monkey cells"** that we wish to have considered for publication in the Proceedings of the National Academy of Sciences (PNAS). The manuscript highlights the efficacy of a protease inhibitor (K777) in blocking SARS CoV-2 infection of several human and primate cell types. The mechanism of action is confirmed as specific inhibition of cathepsin L to prevent host cell processing of the viral spike protein, a requirement for viral cell entry. The inhibitor that is the focus of this manuscript has been approved by the FDA as a clinical-stage compound.

This is a multi-author manuscript with the following **Acknowledgements**:

Funding for experiments completed at Utah State University was provided by the Respiratory Diseases Branch, National Institute for Allergy and Infectious Diseases, NIH USA (Contract N01-AI-30048). Funding for work in the Meek lab at Texas A&M was from AgriLife Research, Texas A&M University. For the Tseng lab, NIAID, Grant #: R24 AI120942 NPARS-S01. Part of the funding for studies performed at UTMB was provided by Selva Therapeutics, Inc.

**There are 25 references and the word count is 4562**

One of the authors, Felix Frueh, is Co-Founder and CSO of Selva Therapeutics. Selva provided partial funding to the Texas Medical Branch for some of the studies reported. James McKerrow is a consultant for Selva Therapeutics. None of the other authors have conflicts of interest related to this manuscript.

**Potential Editorial Board Member**: James Wells

**Potential Members and Editors:** Stephen Beverly
Benjamin Cravat

**Authors and their affiliations:**

Drake M. Mellott,[1] ThChien-Te Tseng,[2,3] Aleksandra Drelich,[3] Jason Hsu,[3] Vivian Tat,[3] Linfeng Li,[1] Frank M. Raushel,[2] Zane Taylor,[2] Ardala Katzfuss,[1] Jiyun Zhu,[1] Miriam A. Giardini,[4] Jair Lage de Siqueira-Neto,[4] Anthony O'Donoghue,[4] Danielle Skinner,[4] Ken Hirata,[4] Pavla Fajtova,[4] Sungjun Beck[4], Alex Clark[4], Laura Berrati[4], Felix Frueh,[5] Brett L. Hurst,[6] Hong Wang,[6] Demetrios Kostomiros[1], Thomas D. Meek[1], James H. McKerrow[4]

**Skaggs School of Pharmacy & Pharmaceutical Sciences**
UC San Diego • 9500 Gilman Drive, # 0657 • La Jolla, CA 92093
T: 858-822-7801 • F: 858-822-5591 • • pharmacy.ucsd.edu

UC San Diego

Skaggs School of Pharmacy
and Pharmaceutical Sciences

Departments of Biochemistry and Biophysics[1] and Chemistry,[2] Texas A&M University, 301 Old Main Drive, College Station, Texas 77843, [3]Department of Microbiology and Immunology, University of Texas, Medical Branch, 3000 University Boulevard, Galveston, Texas, 77755-1001, [4]Skaggs School of Pharmacy and Pharmaceutical Sciences, University of California San Diego, La Jolla, CA, [5]Selva Therapeutics, and [6]Institute for Antiviral Research, Department of Animal, Dairy, and Veterinary Sciences, 5600 Old Main Hill, Utah State University, Logan, Utah, 84322.

Sincerely,

James H. McKerrow, PhD, MD
Dean, Skaggs School of Pharmacy & Pharmaceutical Sciences
Associate Vice Chancellor for Health Sciences

**Skaggs School of Pharmacy & Pharmaceutical Sciences**
UC San Diego • 9500 Gilman Drive, # 0657 • La Jolla, CA 92093
T: 858-822-7801 • F: 858-822-5591 • • pharmacy.ucsd.edu
